# Inhibition of Sting rescues lupus disease by the regulation of Lyn-mediated dendritic cell differentiation

**DOI:** 10.1101/810192

**Authors:** Arthid Thim-uam, Thaneas Prabakaran, Mookmanee Tansakul, Jiradej Makjaroen, Piriya Wongkongkathep, Naphat Chantaravisoot, Thammakorn Saethang, Asada Leelahavanichkul, Thitima Benjachat, Søren Paludan, Trairak Pisitkun, Prapaporn Pisitkun

## Abstract

SLE (systemic lupus erythematosus) is an autoimmune disease that causes chronic inflammation and leads to fatality if left untreated. Immune complex-mediated inflammation and type I IFN signaling pathways are one of the mechanisms initiating lupus disease. Signaling through stimulator of interferon genes (STING) leads to the production of type I IFN and inflammatory cytokines. However, the role of STING in lupus mouse models is controversy. Here we demonstrated the mechanisms of STING involving in SLE pathogenesis at the molecular level. The disruption of STING signaling rescued lupus disease in *Fcgr2b*-deficient mice. STING activated DC facilitated T cell proliferation, which depended on intrinsic expression of STING on DC but not on T cells. Upon STING activation, LYN was recruited and co-localized with STING in bone marrow-derived dendritic cells (BMDC). STING signaling induced phosphorylated LYN and AKT. The inhibition of LYN prohibited STING-induced DC differentiation. Adoptive transfer of STING-activated BMDC into the FCGR2B and STING double-deficiency mice restored lupus phenotypes. These findings provide the proof of concept that inhibition of STING signaling is a promising therapeutic approach for SLE patients.

## Introduction

Systemic lupus erythematosus (SLE) is an autoimmune disease with characteristics of autoantibody production and immune complex deposition that lead to severe inflammation and fatal glomerulonephritis. The heterogeneity of lupus disease has been shown through several mouse models of lupus disease, suggesting a variety of unique mechanisms participating in its pathogenesis. Type I IFN (IFN-I) play significant roles in SLE pathogenesis ^1^. In many SLE patients, the expression of interferon-inducible genes has increased in their peripheral blood mononuclear cells ^2^. Nucleic acid-sensing pathways are the main contributors of IFN-I production ^3^. Several studies suggest that inappropriate recognition of self-nucleic acids can induce the production of IFN-I and promote SLE disease ^1^.

Nucleic acids derived from extracellular sources are sensed via endosomal TLRs, whereas the recognition of cytosolic nucleic acids is independent of TLRs ^4^. The activation of TLRs, such as TLR7 and TLR9, by endosomal nucleic acids, leads to IFN-I production ^5^. Spontaneous duplication of *Tlr7* causes autoimmune lupus phenotypes in *Yaa*-carrying BXSB mice ^6^. Overexpression of *Tlr7* promotes autoimmunity through dendritic cell proliferation, while the deletion of *Tlr7* in lupus-prone MRL/*lpr* mice diminishes autoantibody and immune activation ^7, 8^. However, blocking TLR-mediated signaling by anti-malarial drugs can only treat SLE with mild disease activity ^9, 10^. Thus, investigation of other nucleic acid sensor pathways involved in lupus development could offer a more significant therapeutic opportunity.

Several cytosolic DNA sensors can induce IFN-I production, with cyclic GMP-AMP synthase (cGAS) being the major one ^11^. Cytosolic DNA sensing is also essential for innate immune signaling, and dysregulation of this process can cause autoimmune and inflammatory diseases ^12^. Stimulator of interferon genes (STING), also known as transmembrane protein 173 (TMEM173), is a cytoplasmic adaptor protein that acts downstream of cGAS to enhance IFN-I production ^13^. The loss-of-function mutations in a DNA-specific exonuclease gene *TREX1,* resulting in increased cytosolic DNA levels, are observed in the type I interferonopathies Aicardi-Goutieres syndrome (AGS) and chilblain lupus ^14, 15^. Consistent with these scenarios in human, *Trex1*-deficient mice exhibit fatal inflammation and autoimmunity ^16, 17^. Inhibition of the STING pathway in these mice improves their inflammatory condition and survival ^18^. Moreover, the absence of STING rescues embryonic lethality and arthritis development in another nuclease knockout model, i.e., *DNase II*-deficient mice ^19^. The spontaneous lupus mouse models commonly used to study SLE pathogenesis are MRL/*lpr*, NZBxNZW.F1, and BxSB ^20^. Since these models possess different genetic backgrounds, each model could develop lupus with unique pathogenesis ^6, 20, 21^. Surprisingly, the absence of STING in MRL/*lpr* mice does not improve lupus phenotypes but instead promotes more inflammation ^22^. Furthermore, knocking down the IFN receptor gene *Ifnar1* in MRL/*lpr* mice aggravates lymphoproliferation, autoantibody production, and end-organ damage ^23, 24^. These data suggest that the role of STING in lupus pathogenesis is controversy dependent on the models studied. Therefore further studies in a relevant animal model that reflect human lupus are required to circumvent these conflicting data.

A comprehensive genetic analysis has identified *FCGR2B* as a susceptibility gene in SLE patients ^25^. The deletion of the *Fcgr2b* gene causes a lupus-like disease in genetic susceptibility to autoimmune development. The *Fcgr2b^-/-^* mice created in 129 strain with subsequently backcrossed into C57BL/6 develop overt lupus disease ^26, 27^. The 129-derived Sle16 covering the Nba2 interval region is an autoimmune susceptibility locus, which contains the *Fcgr2b*, Slam family, interferon-inducible Ifi200 family genes ^27–29^. Among the Ifi200 family, the *Ifi202* shows the highest expression in the splenocytes from Nba2 carrying mice ^30^. The *Ifi202* is a candidate lupus susceptibility gene, and its human homolog *IFI16* shows the association with SLE ^31^. Also, IFI16 signals through STING to initiate IFN-I production ^32, 33^. Based on the genetic background of the 129-derived locus, STING may play a significant role in the pathogenesis of the 129/B6.*Fcgr2b^-/-^* lupus mice.

In this work, we showed an increase of *Ifi202* and *Sting* expression in the 129/B6.*Fcgr2b^-/-^* mice. Disruption of STING signaling rescued lupus phenotypes of the 129/B6.*Fcgr2b^-/-^* mice. Stimulation of STING promoted dendritic cell maturation and plasmacytoid dendritic cell differentiation. After STING activation, LYN was phosphorylated and recruited to interact with STING. Inhibition of LYN diminished STING driven differentiation of dendritic cells. The adoptive transfer of STING-activated bone marrow-derived dendritic cells (BMDC) into the double-deficiency (*Fcgr2b^-/-^. Sting^gt/gt^*) mice restored the lupus phenotypes. The data suggested that STING signaling in the dendritic cells initiated the autoimmune development in the 129/B6.*Fcgr2b^-/-^* mice. STING is a promising therapeutic target for lupus disease.

## Results

### Loss of the stimulator of type I interferon genes (STING) increases survival of *Fcgr2b^-/-^* lupus mice

First, we confirmed that 129/B6.*Fcgr2b^-/-^* mice (or *Fcgr2b^-/-^* in short) showed the increase of *Ifi202* expression (Fig. 1A). The mRNA and protein expressions of *Sting* in the spleen were also upregulated in the *Fcgr2b^-/-^* mice (Fig. 1B and 1C). We further observed the significant rise of mRNA expression of *Ifnb*, interferon-inducible genes (*Irf7*, *Cxcl10*, and *Mx1*), and *Ifng* in the spleen of the *Fcgr2b^-/-^* mice (Fig. 1D-1H). To determine whether the Sting signaling is required for lupus development in the *Fcgr2b^-/-^* mice, we generated the double deficiency of *Fcgr2b* and *Sting* together with control littermates. The *Fcgr2b^-/-^* mice were crossed with the C57BL/6.*Sting* deficiency or Goldenticket mice (*Sting^gt^*) which behave as a functional knockout of STING ^34^. The double-deficient mice showed a higher survival rate compared to the *Fcgr2b^-/-^* mice (Fig. 1I) while the *Fcgr2b^-/-^* with heterozygote of Sting (*Sting^wt/gt^*) did not alter the phenotypes of *Fcgr2b^-/-^* with wild-type Sting (*Sting^wt/wt^*).

**Figure 1.**
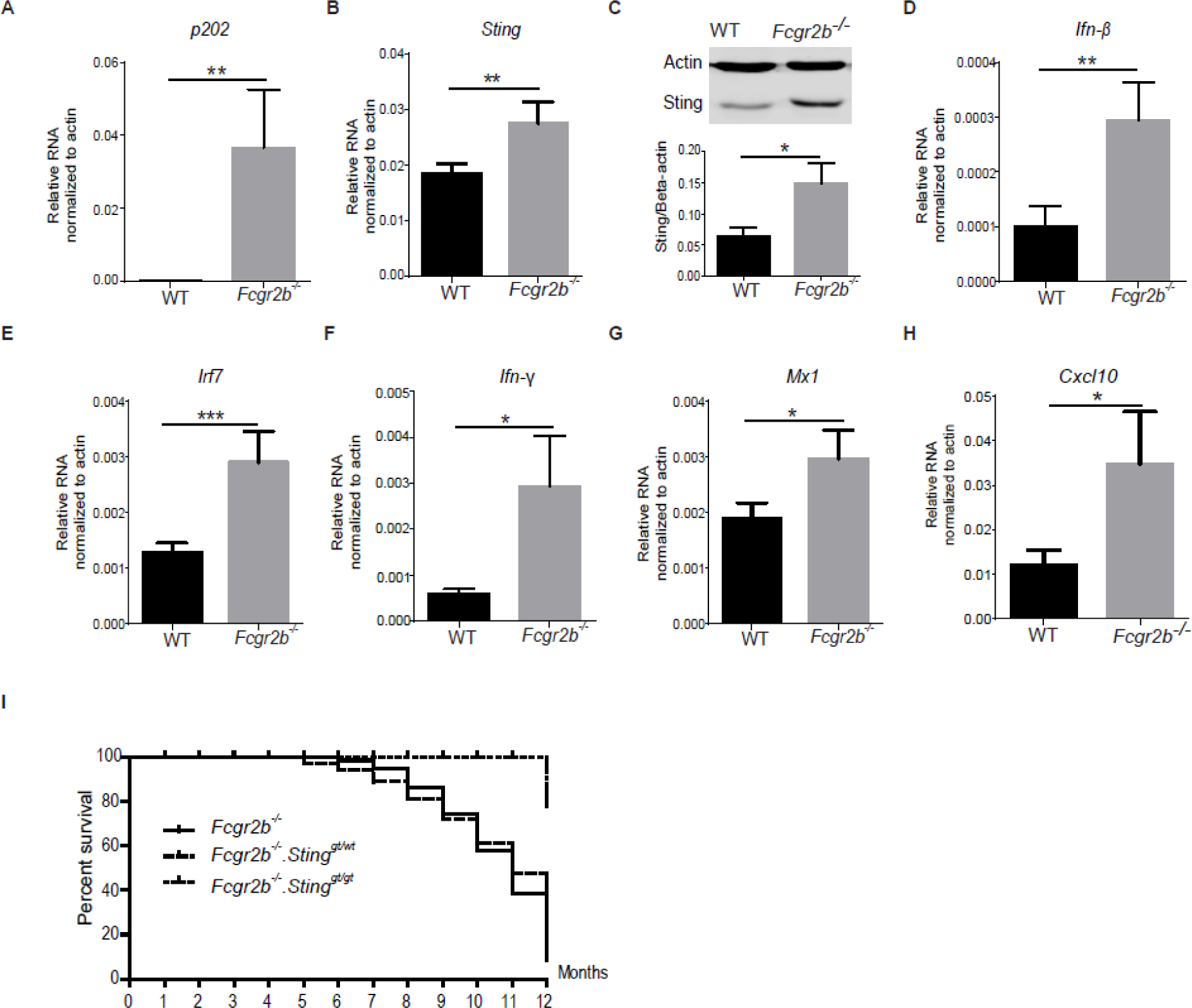
Loss of the stimulator of type I interferon genes (STING) increases the survival of *Fcgr2b^-/-^* lupus mice. Gene expression profiles from spleens of wild-type and *Fcgr2b^-/-^* mice at the age of 6 months were tested by real-time PCR (N=10-12 per group). The mRNA expressions of (A) *p202*, (B) *Sting*, (D) *Ifnβ*, (E) *Irf3*, (F) *Irf7*, (G) *Ifnγ*, (H) *Mx1*, and (I) *Cxcl10* were shown. (C) Isolated splenocytes were analyzed for STING protein expression by western blot. Data are representative of 3 mice per group. Quantification of the intensity was normalized by actin (N=3 per group). (J) The *Fcgr2b*-deficient mice were crossed with *Sting*-deficient mice (*Sting^gt/gt^*) to generate the double-deficient mice (*Fcgr2b^-/-^. Sting^gt/gt^*) and littermate controls. The survival curve of the mice was observed for up to 12 months (N=14 per group). Error bars indicate SEM, *p < 0.05, **p<0.01, and ***p<0.001.

### STING signaling pathway promotes autoantibody production and glomerulonephritis in the *Fcgr2b*^-/-^ lupus mice

The lupus phenotypes of the double-deficient mice were examined and compared with littermate controls. The level of the anti-nuclear antibody (ANA) and anti-dsDNA antibody from the sera of the double-deficient mice (*Fcgr2b^-/-^. Sting^gt/gt^*) showed significantly lower than the *Fcgr2b*^-/-^ mice (Fig. 2A-2C). The kidneys of *Fcgr2b^-/-^* mice showed pathology of diffuse proliferative glomerulonephritis, which did not present in the *Fcgr2b^-/-^*.*Sting^gt/gt^* mice (Fig. 2D). The glomerular and interstitial scores in the kidneys of *Fcgr2b^-/-^* mice were significantly higher than the double-deficient mice (Fig. 2E-2F). Consistent with the pathology, the immunofluorescence staining showed fewer CD45^+^ cells and IgG deposition in the kidneys of *Fcgr2b^-/-^*.*Sting^gt/gt^* mice (Fig. 2G).

**Figure 2.**
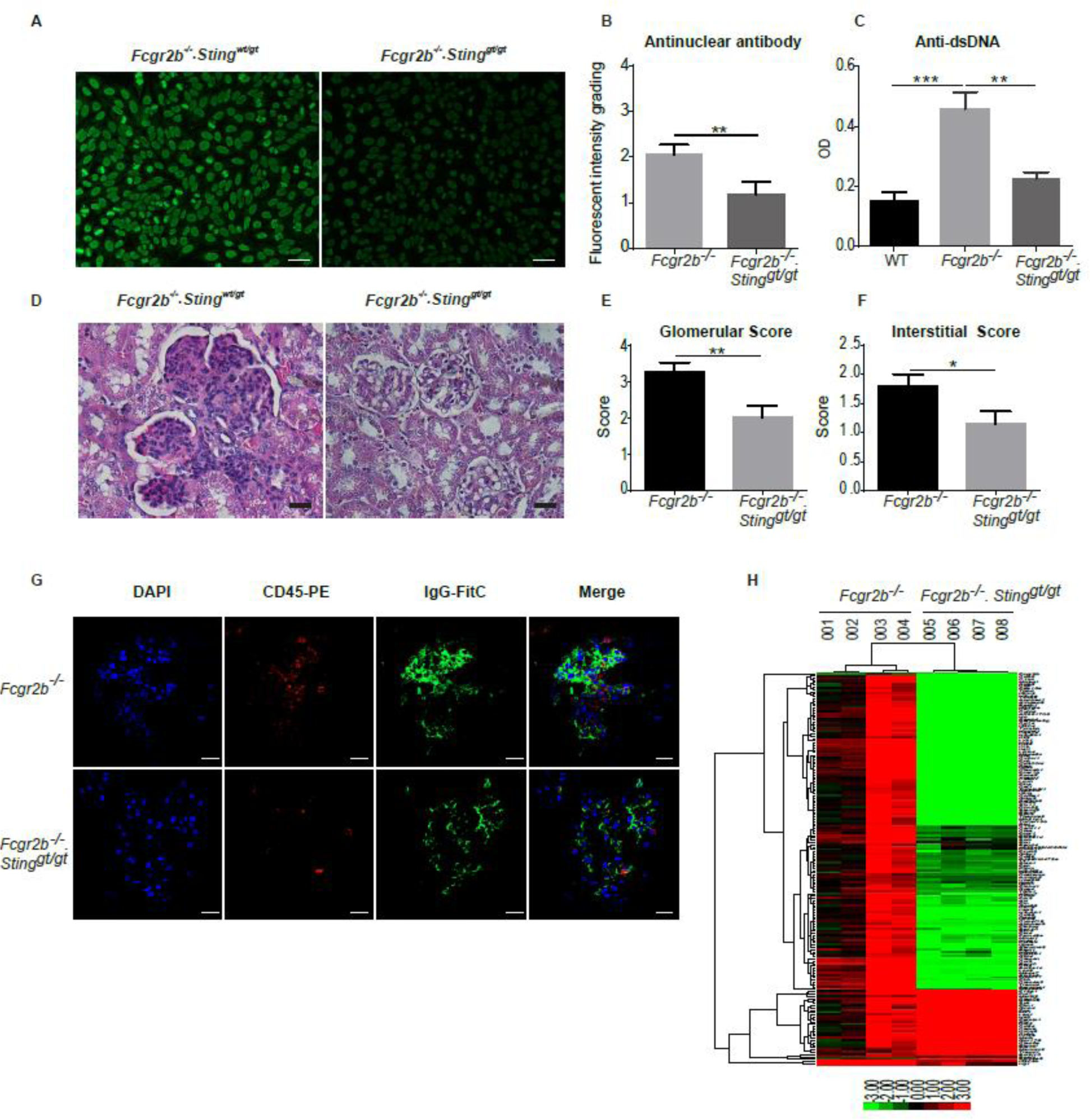
STING signaling pathway promotes autoantibody production and glomerulonephritis in the *Fcgr2b^-/-^* lupus mice. The anti-nuclear antibodies (ANA) from the sera of *Fcgr2b^-/-^* and *Fcgr2b^-/-^. Sting^gt/gt^* (dilution 1:800) show by (A) the immunofluorescence staining. Data are representative of 8 mice per group (scale bar = 20 µm). (B) Semi-quantification of ANA was graded by fluorescence intensity (N=8 mice per group). (C) Anti-dsDNA from sera (dilution 1:100) of *Fcgr2b^-/-^* and *Fcgr2b^-/-^. Sting^gt/gt^* was detected by ELISA (N=10-11 per group). (D) Kidney sections of *Fcgr2b^-/-^* and *Fcgr2b^-/-^. Sting^gt/gt^* mice (6-8 months old) were stained with H&E. Data are representative of 7-10 mice per group (scale bar = 25 µm). (E-F) Glomerular scores and interstitial scores of kidney sections were blindly graded (N=7–10 per group). Data show as mean ± SEM (*p < 0.05, **p<0.01 and ***p<0.001). (G) Immunofluorescence stainings of the kidneys from *Fcgr2b^-/-^* and *Fcgr2b^-/-^. Sting^gt/gt^* mice show in green (IgG), red (CD45), and blue (DAPI). Data are representative of 3 mice per group (scale bar=10 µm). (H) A heat map of microarray data from the kidneys of *Fcgr2b^-/-^* and *Fcgr2b^-/-^. Sting^gt/gt^* mice show that the interferon signature genes significantly changed in the *Fcgr2b^-/-^*mice (N=4 mice per group). Data show in log2 (sample/wild-type).

We further looked at the gene expression profiles in the kidney of these mice and found a significantly different expression (Fig. S1A). The expression of interferon-inducible genes in the kidney of these mice was higher in the *Fcgr2b^-/-^* mice, especially in the ones with greater severity (#003, 004) and a significant reduction of interferon-inducible genes in the kidneys of *Fcgr2b^-/-^*.*Sting^gt/gt^* mice (Fig. 2H). However, not all of these interferon-inducible genes decreased in the *Fcgr2b^-/-^*.*Sting^gt/gt^* mice (Fig. 2H). This data suggested that STING-dependent pathology mediated by both interferon and non-interferon signaling and not all of the interferon signaling in the *Fcgr2b^-/-^* mice contributed by the STING pathway.

### STING is essential for inflammatory phenotypes of the *Fcgr2b*^-/-^ lupus mice

The expression of interferon-inducible genes in the kidneys was confirmed by real-time PCR. The expressions of *Isg15*, *Irf7*, and *Mx1* were upregulated in the *Fcgr2b^-/-^* mice and downregulated in the absence of STING (Fig. 3A-3C). Also, the expression of *Irf5*, the lupus susceptibility gene, that upregulated in the kidneys of the *Fcgr2b^-/-^* mice was Sting-dependent (Fig. 3D). The splenocytes were analyzed from the mice at the age of 6-7 months to characterize the immunophenotypes. The expansion of dendritic cells (CD11c^+^) and plasmacytoid dendritic cells (CD11c^+^PDCA^+^) in the *Fcgr2b^-/-^* mice diminished in the absence of Sting (Fig. 3E-3F). The reduction of T effector memory cells (CD3^+^CD4^+^CD62L^lo^CD44^hi^), germinal center B cells (B220^+^GL7^+^), and CD3^+^CD4^+^ICOS^+^ cells in the double-deficient mice (Fig. 3G-3H, and Fig. S2A). Also, the mean fluorescence intensity of MHC-II (IA-b) on B cells significantly reduced in the double-deficient mice (Fig. S2B). However, the expansion of plasma cells did not show the difference between single and double-deficient mice (Fig. S2C). Furthermore, the sera levels of MCP-1 and TNF-α from the *Fcgr2b^-/-^* mice were significantly increased compared to WT mice (Fig. 3I-3J), whereas IL-1β and IL-23 did not show the significant changes (Fig. 3K-3I). However, the levels of TNF-α, IL-1β, and IL-23 from the *Fcgr2b^-/-^* mice significantly decreased in the absence of STING (Fig. 3J-3I). These data suggested that STING mediated the inflammatory process in the *Fcgr2b*-deficient lupus mice.

**Figure 3.**
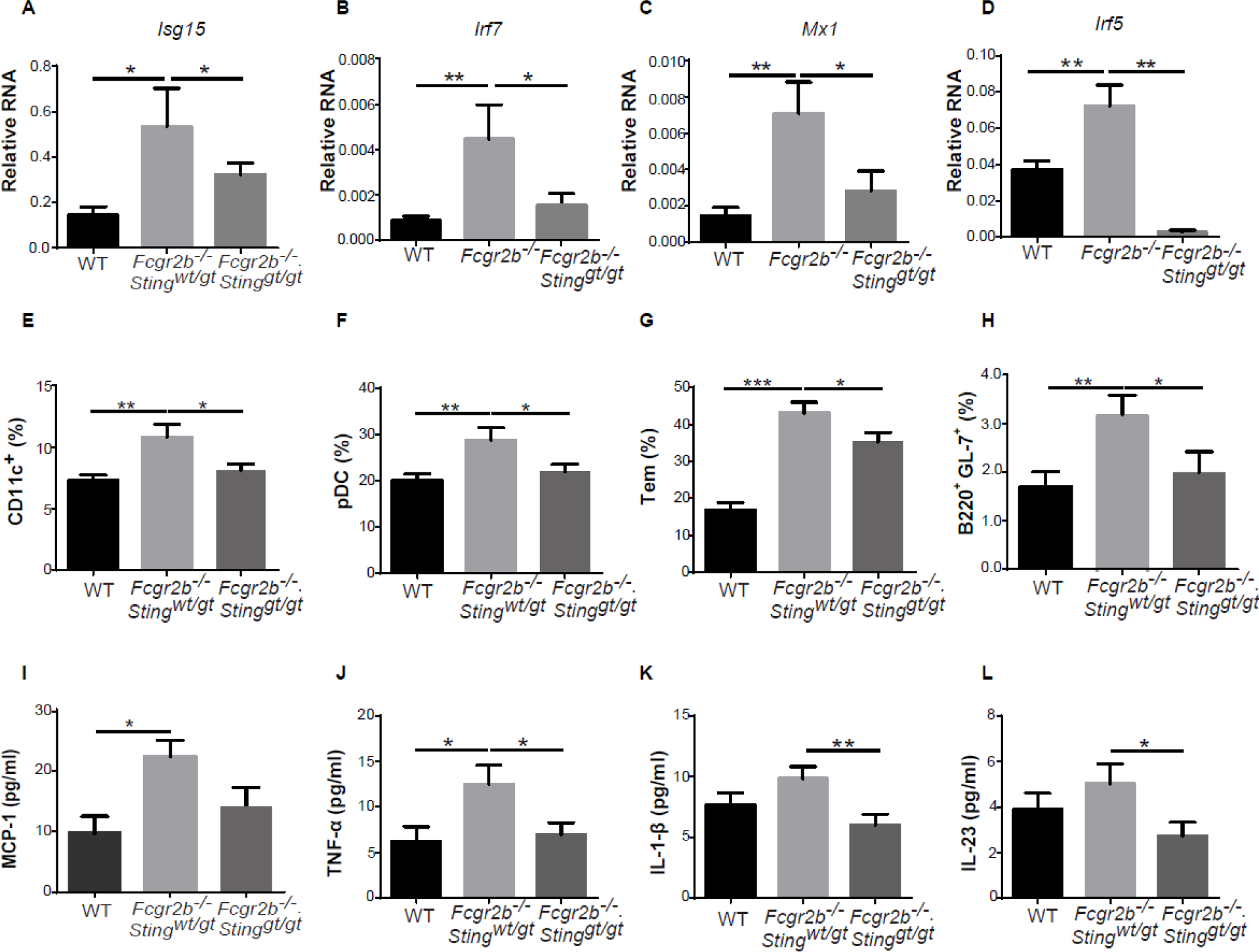
STING is essential for inflammatory phenotypes of the *Fcgr2b^-/-^* lupus mice. (A-D) Gene expression profiles from the kidneys of wild-type, *Fcgr2b^-/-^* and *Fcgr2b^-/-^. Sting^gt/gt^* mice at the age of 6 months were tested by real-time PCR (N=10-17 per group). The mRNA expressions of interferon-inducible genes shown in (A) *Irf3*, (B) *Irf5*, (C) *Irf7*, and (D) *Mx1*. (E-H) Flow cytometry analysis of splenocytes isolated from wild-type, *Fcgr2b^-/-,^* and *Fcgr2b^-/-^. Sting^gt/gt^* mice at the age of 6-7 months (N= 13-14 per group). Data shown in the percentage of (E) CD11c^+^, (F) plasmacytoid dendritic cells (pDC), (G) Tem (CD3^+^CD4^+^CD44^hi^CD62L^lo^), and (H) B220^+^GL7^+^ cells. (I-L) The sera cytokines of wild-type, *Fcgr2b^-/-^* and *Fcgr2b^-/-^. Sting^gt/gt^* mice at the age of 6 months were analyzed by cytometric bead array. Serum cytokines of (I) MCP-1, (J) TNF-α, (K) IL-1β, and (L) IL-23 (N=10-15 per group). Data show as mean ± SEM (*p < 0.05, **p<0.01 and ***p<0.001).

### STING-activated dendritic cells induce the proliferation of naïve CD4*^+^* T cells

The splenocytes of *Fcgr2b^-/-^* mice showed an increase of effector memory T cells (Tem) and *Ifng* expression (Fig. 3G and 1F). We hypothesized that the high proportion of Tem in the *Fcgr2b^-/-^* mice might contribute to the increase of *Ifng* expression. The intercellular staining of IFN-γ was performed to test this assumption. The IFN-γ producing CD4^+^ T cells from the *Fcgr2b^-/-^* mice were higher than wild-type and double-deficient mice (Fig. 4A-4C). The flow cytometry data showed that the expansion of DC in the spleen of the *Fcgr2b^-/-^* mice was STING-dependent (Fig. 3E). T cells were co-cultured with BMDC to check if the *Sting*-expressing DC could directly influence the T cell phenotypes. The short-interval (6 hours) co-cultured between STING-activated BMDC derived from *Fcgr2b^-/-^* and double-deficient mice and CD4^+^ T cells showed comparable numbers of IFN-γ producing CD4^+^ T cells which were independent of *Sting* expression on BMDC (Fig. 4D-4E). However, the CD4^+^ T cells derived from the *Fcgr2b^-/-^* mice expressed the intracellular staining of IFN-γ higher than the double-knockout mice when co-cultured with *Sting*-sufficient BMDC (Fig. 4D-4E).

**Figure 4.**
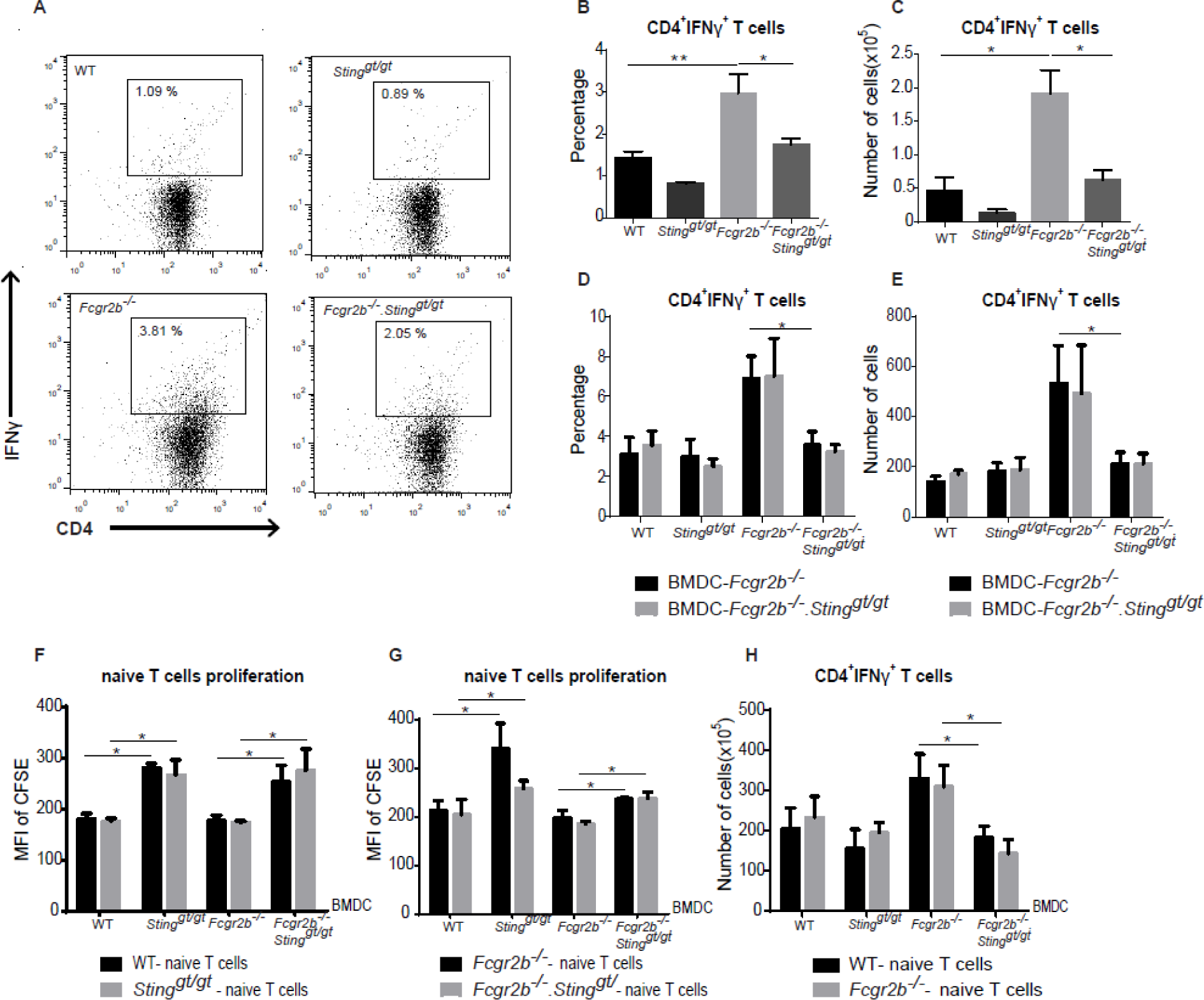
STING-activated dendritic cells induce the proliferation of naïve CD4+ T cells. (A-C) Flow cytometry analysis of (A-B) intracellular staining of IFN-γ -producing CD4^+^ T cells isolated from lymph nodes of wild-type, *Sting^gt/gt^*, *Fcgr2b^-/-,^* and *Fcgr2b^-/-^. Sting^gt/gt^* mice at the age of 6-7 months. (A) Data are representative of 3-4 mice per group. (B) The percentage of IFN-γ^+^CD4^+^ T cells and (C) the number of IFN-γ^+^CD4^+^ T cells (N=3-4 per group). (D) The percentage and (E) the number of intracellular IFN-γ producing CD4^+^ cells after co-cultured with DMXAA activated BMDC from *Fcgr2b^-/-^* and *Fcgr2b^-/-^. Sting^gt/gt^* (6–7 months old) for six hours (N=3-4). (F-H) Co-culture of naïve T cells with DMXAA activated BMDC from wild-type, *Sting^gt/gt^*, *Fcgr2b^-/-^*, and *Fcgr2b^-/-^. Sting^gt/gt^* mice (labeled at x-axis) for 72 hours. (E and G) CFSE dilution of isolated naïve T cells showed in mean fluorescence intensity (MFI), and (H) the total numbers of IFN-γ^+^CD4^+^ T cells (N=4 per group). Data show as mean ± SEM (*p < 0.05, and **p<0.01).

Next, we examined if Sting activated BMDC can promote T cell proliferation. The co-cultured of the Sting-activated BMDC with purified naïve T cells for 72 hours was assessed with CFSE dilution assay (Fig. 4F-4G). The proliferation ability of naïve T cells was independent of *Sting* expression on T cells but depended on *Sting*-expression on BMDC (Fig. 4F-4G). The naïve T cells that primed with *Sting* expressing BMDC derived from the *Fcgr2b^-/-^* mice were capable of producing higher IFN-γ compared to the cells that were co-cultured with double-deficient BMDC (Fig. 4H). These data suggest that STING activation promoted the maturation of DC, which subsequently primed the naïve CD4^+^ T cells to proliferate and become the IFN-γ producing T cells.

### STING activation promotes the maturation of dendritic cells and the differentiation of plasmacytoid dendritic cells

The cGAS/STING pathway is essential for DC activation ^35^. STING activating DC can induce naïve T cells to proliferate and produce IFN-γ (Fig. 4F-4H), which suggested STING may enhance the maturation of DC to become professional antigen-presenting cells (APC). To test if the expansion of DC in the *Fcgr2b^-/-^* mice is directly mediated by STING signaling, the bone marrow-derived dendritic cells (BMDC) were differentiated into immature DC and subsequently stimulated with STING ligands (DMXAA), DMSO, and LPS (as a control) to assess if STING played a role in DC maturation. The LPS control induced the immature DC to increase the expression of MHC-II (IA-b) and CD80, which suggested the phenotypes of mature DC, from both *Sting*-sufficient and *Sting*-deficient mice (Fig. 5A and 5C). The immature DC from wild-type and *Fcgr2b^-/-^* mice also showed the increasing percentage of IA-b^+^ and CD80^+^ DC cells after DMXAA stimulation; the *Sting*-deficient mice did not develop these mature phenotypes (Fig. 5B and 5D). The supernatant from BMDC culture with DMXAA stimulation showed an increase in the concentration of IL-1α, IL-6, TNF-α, and MCP-1 in the wild-type and *Fcgr2b^-/-^* mice but not in *Sting*-deficient mice and double-deficient mice (Fig. 5E-5H).

**Figure 5.**
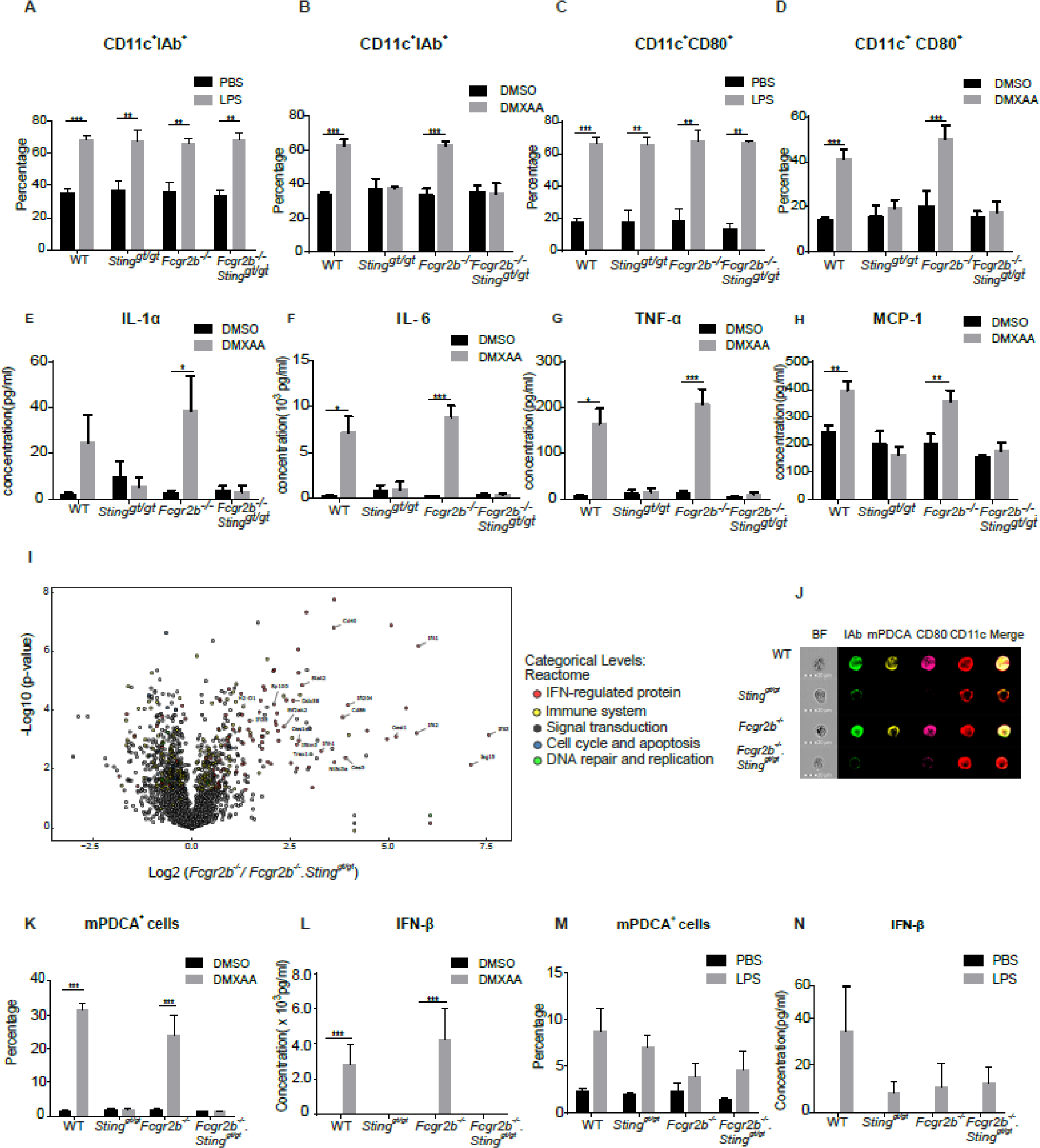
STING activation promotes the maturation of dendritic cells and the differentiation of plasmacytoid dendritic cells. Bone marrows were isolated from wild-type, *Sting^gt/gt^*, *Fcgr2b^-/-^* and *Fcgr2b^-/-^. Sting^gt/gt^* mice at the age of 6 months. (A-D) IL-4 and G-CSF differentiated bone marrow-derived dendritic cells (BMDC) for five days were stimulated with LPS or DMXAA for 24 hours. Flow cytometry analysis shows the percentage of (A-B) CD11c^+^ IAb^+^ cells and (C-D) CD11c^+^CD80^+^ cells. (E-H) Supernatants were collected and analyzed after DMXAA stimulation for 24 hours. Cytometric bead array shows the levels of (E) IL-1α, (F) IL-6, (G) TNF-α, and (H) MCP-1. (I) Volcano plot of protein expressions from proteomic analysis of DMXAA activated BMDC of *Fcgr2b^-/-^* and *Fcgr2b^-/-^*.*Sting^gt/gt^* mice at the age of 6-7 months (N=4 per group). (J) Imaging flow cytometry of DMXAA activated BMDC shows the representative staining of IAb (green), mPDCA (yellow), CD80 (pink), and CD11c (red) (N= 3 mice per group). (K and M) The percentage of pDC (PDCA^+^ cells) after (K) DMXAA activation and (M) LPS activation for 24 hours (N = 3–4 per group). (L and N) The level of IFN-β from the culture supernatant of activated BMDC with (L) DMXAA and (N) LPS (N = 5 per group). Data show as mean ± SEM (N=3–5; *p < 0.05, **p<0.01 and ***p<0.001).

To better understand the mechanisms of STING in DC differentiation, we performed the proteomic analysis of STING-activated BMDC in the *Fcgr2b^-/-^* mice compared to the double-deficient mice. The Volcano plot showed the protein that highly expressed were interferon-regulated proteins (Fig. 5I and Table S1). This finding may result from the increase of IFN-I production in the culture medium, which could upregulate the interferon-regulated proteins. We hypothesized that STING might promote the differentiation of pDC (a major producer of IFN-I). The in vitro culture of BMDC with DMXAA and LPS (as a control) showed a significant increase in pDC and IFN-β with DMXAA but not with LPS stimulation (Fig. 5K-5N). Also, we demonstrated the morphology of these cells by the imaging flow cytometry and found the pDC expressed CD80 as well (Fig. 5J). These data suggested that Sting signaling pathway mediated DC maturation and pDC differentiation.

### STING mediated maturation of conventional DC and differentiation of plasmacytoid DC via LYN signaling pathway

STING-interacting proteins were identified by immunoprecipitation (IP) using the STING antibody that targets the N-terminal region of the protein. STING-activating BMDC with DMXAA for 3 hours was immunoprecipitated and analyzed by mass spectrometry (Table S2). Among the proteins detected, LYN, a member of Src family kinases (SFK), has been shown its function in the maturation of DC and pDC response ^36, 37^. Immunoprecipitation confirmed that LYN interacted with STING after DMXAA stimulation in WT BMDC while LYN constitutively interacted with STING in the *Fcgr2b*-deficient BMDC (Fig. 6A). This interaction could result from the intrinsic activation of STING in the *Fcgr2b^-/-^* mice. This activation did not change the protein abundance of LYN and STING in total cell lysate (Fig. 6B). Also, the western blot from both IP and cell lysate showed another fainting band of STING after the activation; this protein was identified by mass spectrometry as the phosphorylation of STING (Ser357) (Fig. S3A-S3B).

**Figure 6.**
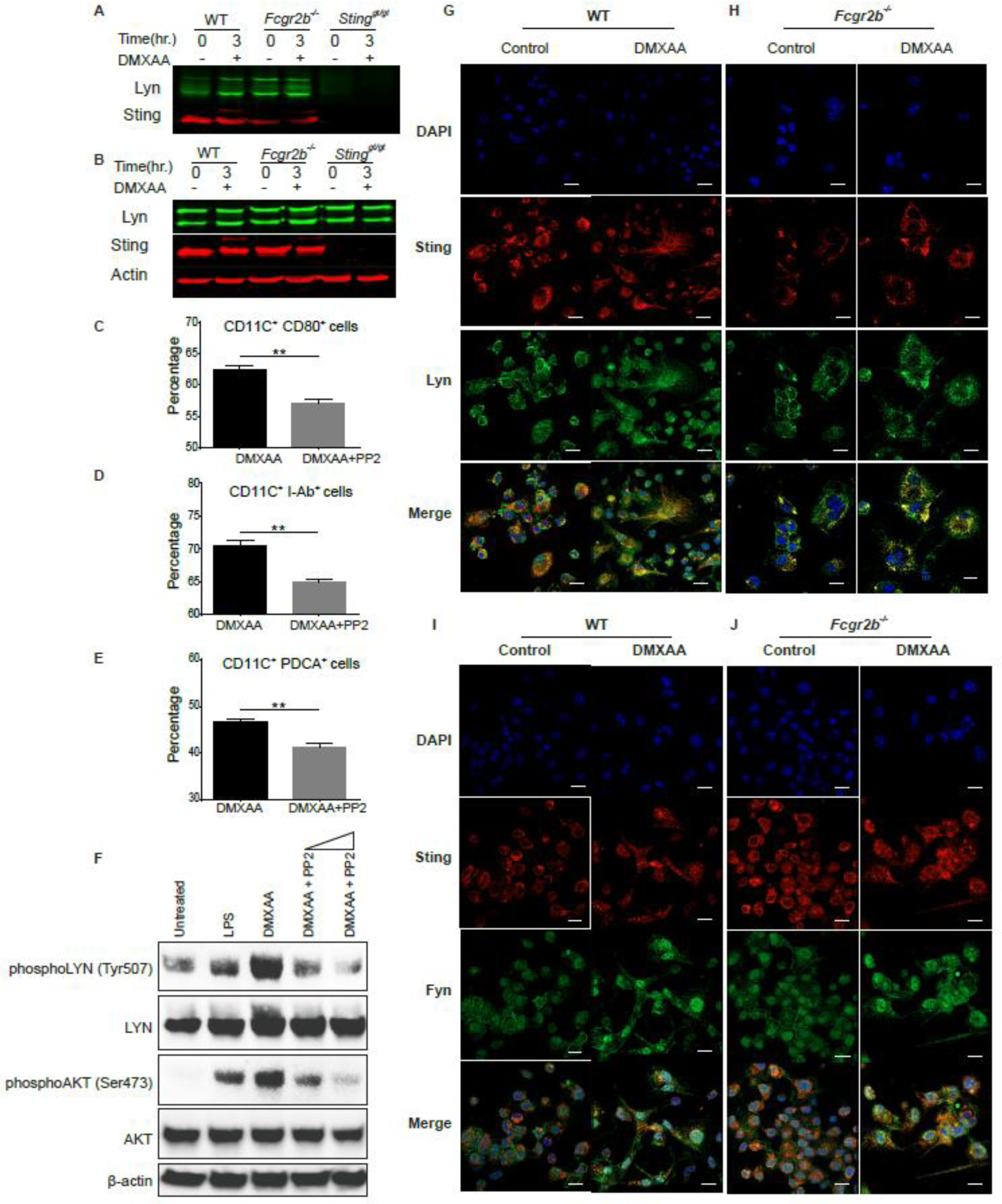
STING mediated maturation of conventional DC and differentiation of plasmacytoid DC via LYN signaling pathway. (A and B) Fluorescent western blot shows (A) the immunoprecipitation with STING (red) and blots with LYN (green) and (B) cell lysate of activated BMDC with DMXAA at 0, and 3 hr. A representative of 4 experiments. STING-activated BMDC were co-cultured with LYN inhibitor PP2 and analyzed by (C-E) flow cytometry, which showed the percentage of (C) CD80^+^CD11c^+^, (D) I-Ab^+^CD11c^+^ and (E) PDCA^+^CD11c^+^ cells (N = 3 per group). (F) Western blot analysis of STING-activated BMDC with or without PP2 inhibitor showed the phosphorylation of LYN (Try507) and AKT (Ser473). Data are representative of 3 mice per group. (G-J) The confocal microscope of DMXAA activated BMDC from WT and *Fcgr2b^-/-^* mice for 6 hours. (G and H) Staining BMDC shows LYN (green), STING (red), and DAPI (blue). (I and J) Staining BMDC shows FYN (green), STING (red), and DAPI (blue) (scale bar=20 um). The representative of 5 experiments. Data show as mean± SEM (*p < 0.05, **p<0.01, and ***p<0.001).

Lyn kinase regulates DC maturation, and genetic deletion of *Lyn* ablates pDC ^36, 37^. We hypothesized that STING promoted DC maturation and pDC differentiation through LYN signaling pathway, and the inhibition of LYN should affect the STING-induced BMDC differentiation. Our in vitro data showed that pan SFK inhibitor PP2 decreased STING-mediated expression of IAb and CD80 on conventional DC (Fig. 6C-6D and Fig. S3C) and the differentiation of pDC (Fig. 6E). The activation of STING increased the phosphorylation of LYN (Tyr507) and AKT (Ser473), which were inhibited by PP2 (Fig. 6F). However, FYN, a member of the Src family kinases (SFK), has been shown the functional role in pDC maturation and PP2 inhibited both LYN and FYN signaling ^37^. Therefore, we identified the colocalization of LYN, FYN, and STING to confirm the protein interaction, and found that LYN, but not FYN, showed the colocalization with STING in the STING-activated BMDC (Fig. 6G-6J). The *Fcgr2b*-deficient BMDC constitutively showed a certain degree of the colocalization between STING and LYN, while the activation of STING promoted more interaction (Fig. 6H). The data suggested that STING mediated maturation and differentiation of conventional DCs and pDCs, respectively, is dependent on LYN signaling pathway.

### STING activated BMDC induce autoimmunity in the WT mice

The STING signaling pathway activated the immature BMDC to differentiate into the mature DC and pDC. Dendritic cells are significant producers of inflammatory cytokines and capable of promoting T cell proliferation and differentiation. We proposed that STING may induce the lupus disease by initially acting through the DC activation. We performed the adoptive transfer of the STING-activated BMDC derived from *Fcgr2b^-/-^* mice into WT recipient mice to test this hypothesis. The recipient WT mice developed the autoimmune phenotypes, including the production of anti-dsDNA (Fig. 7A), expansion of Tem, CD4+ICOS+ cells, plasma cells, and germinal center B cells (Fig. 7B-E). The STING-activated BMDC from *Fcgr2b^-/-^* mice also induced the immune complex deposition in the kidneys of recipient WT mice (Fig. 7F). These data suggested that the STING-activated BMDC of *Fcgr2b^-/-^* mice promoted autoimmunity conditions.

**Figure 7.**
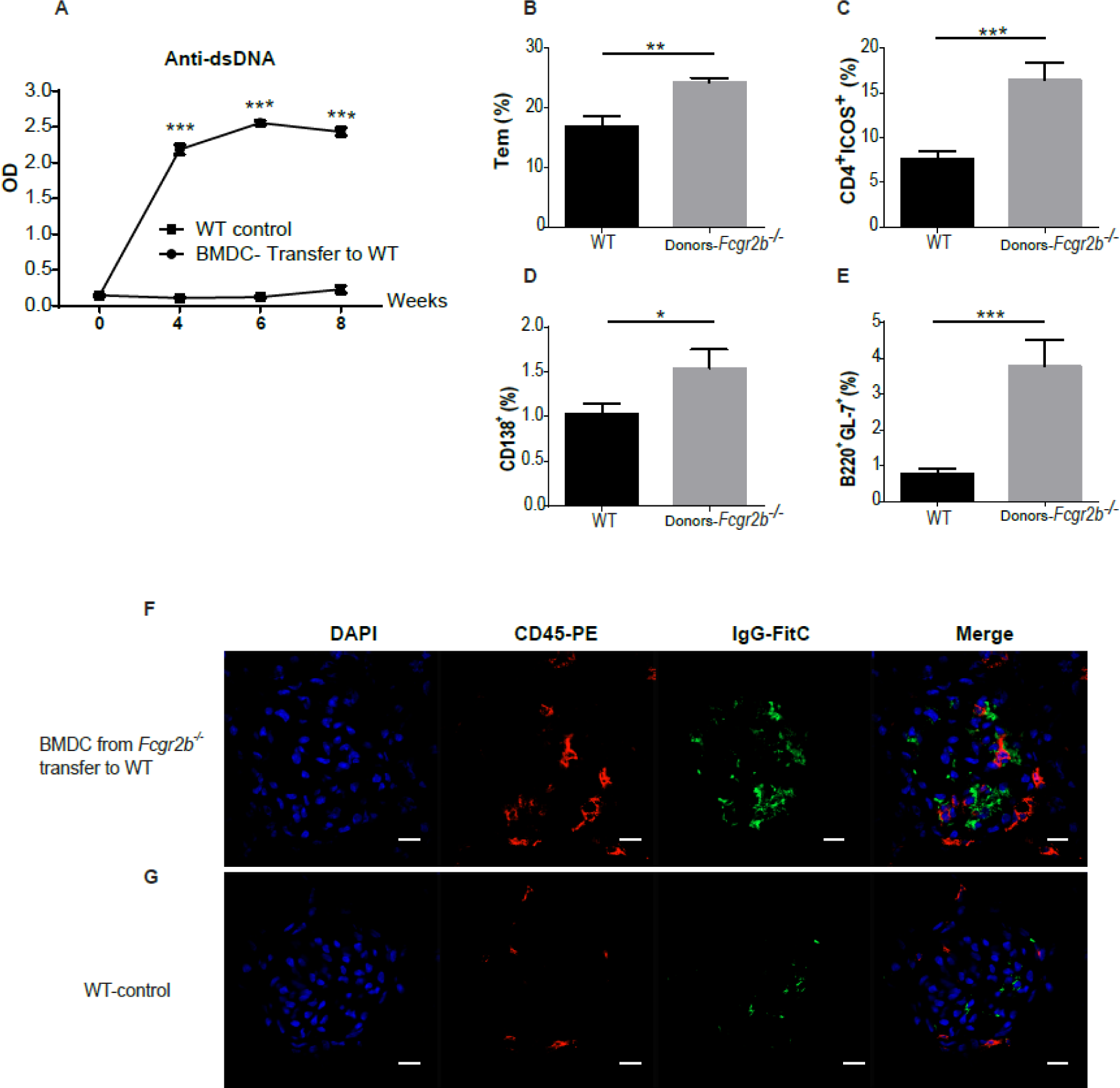
STING activated BMDC induce autoimmunity in WT mice. (A-G) DMXAA-activated BMDC from *Fcgr2b^-/-^* mice were transferred into the WT recipient mice. (A) The level of anti-dsDNA from the sera (1:100) measured by ELISA (N=4-5 per group). Flow cytometry analysis of splenocytes from the recipient after BMDC transferred every two weeks for four times shows the percentage of (B) effector T cells (CD4^+^CD44^hi^CD62L^lo^), (C) CD4^+^ICOS^+^ cells, (D) CD138^+^ cells and (E) B220^+^GL7^+^ cells (N=4-7 per group). Data show as mean ± SEM (*p < 0.05, **p<0.01 and ***p<0.001). (F and G) Immunofluorescence stainings of the kidneys from recipient wild-type mice show in green (IgG), red (CD45), and blue (DAPI). Data are representative of 4 mice per group (scale bar=10 μm).

### Sting expressing BMDC induce lupus development in the *Fcgr2b^-/-^. Sting ^gt/gt^* mice

Next, we performed the reconstitution experiment by adoptive transfer of STING-activated BMDC into the double-deficient mice. The level of anti-dsDNA significantly increased in the recipient mice of *Sting*-sufficient BMDC compared to receiving *Sting*-deficient BMDC and controlled (Fig. 8A).

**Figure 8.**
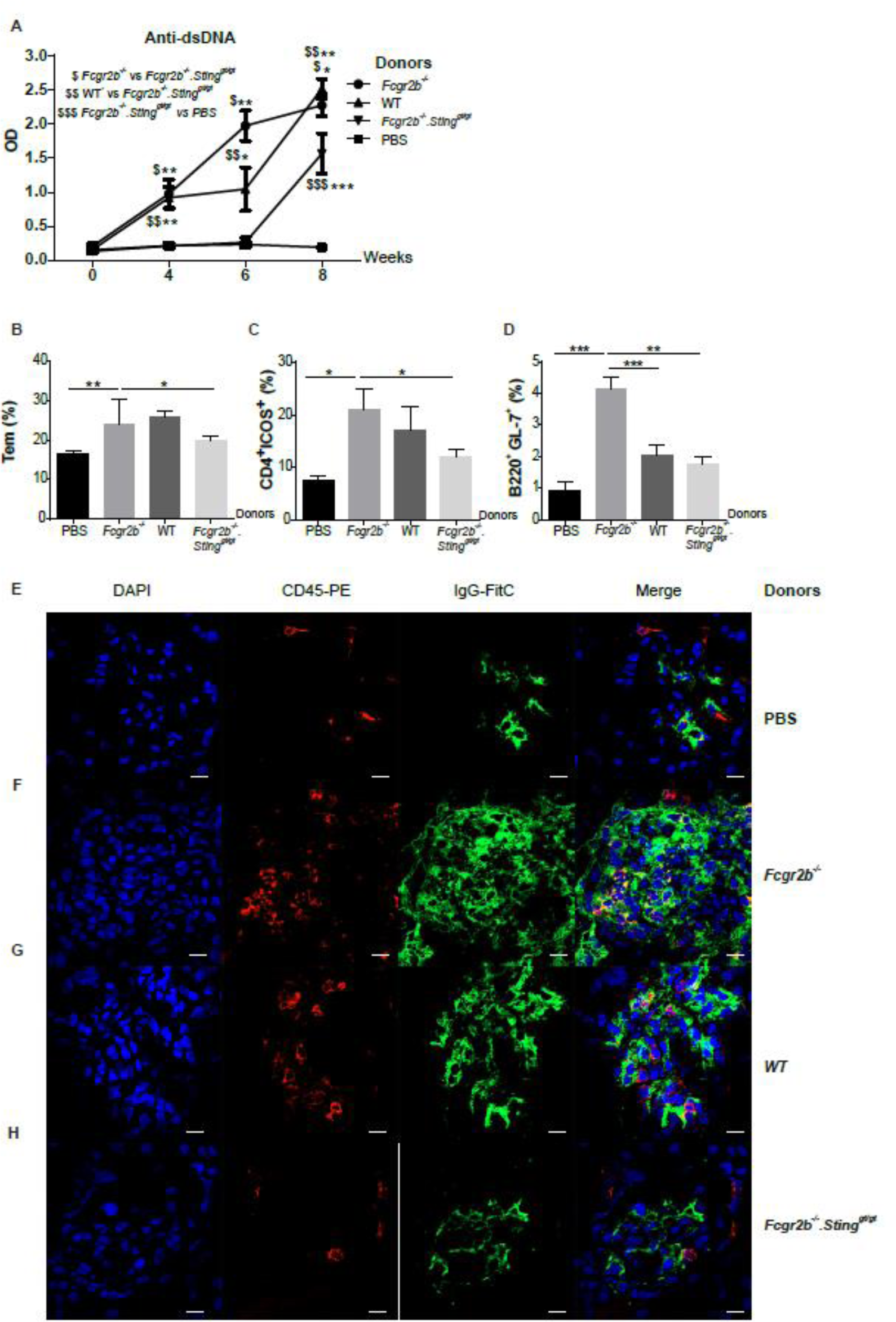
Sting expressing BMDC induce lupus development in the *Fcgr2b^-/-^. Sting ^gt/gt^* mice. **(A-H)** DMXAA activated BMDC from *Fcgr2b^-/-^*, WT, and *Fcgr2b^-/-^*.*Sting^gt/gt^* were transferred into the recipient mice (*Fcgr2b^-/-^. Sting^gt/gt^*). (A) The level of anti-dsDNA from the sera (1:100) measured by ELISA (N=5-10 per group). (B-E) Flow cytometry analysis of recipient splenocytes after BMDC transferred every two weeks for four times shows the percentage of (B) Tem (CD4^+^CD44^hi^CD62L^lo^), (C) CD4^+^ICOS^+^ cells, (D) B220^+^GL7^+^ cells and (E) B220^+^IAb^+^ cells (N=5-10 per group). (F-I) Immunofluorescence staining of the kidney from the *Fcgr2b^-/-^*.*Sting^gt/gt^* recipient mice after the transfer with (F) PBS control, DMXAA activated BMDC from (G) *Fcgr2b^-/-^*, (H) WT and (I) *Fcgr2b^-/-^. Sting^gt/gt^*. The confocal microscope shows DAPI (blue), CD45 (red), and IgG (green). The representative of 3 experiments (scale bar=10 um). Data show as mean ± SEM (*p < 0.05, **p<0.01, and ***p<0.001).

The flow cytometry analysis of spleens from recipient mice receiving either STING-activated BMDC derived from WT or *Fcgr2b*^-/-^ mice showed the increase in the percentage of Tem, and CD4^+^ICOS^+^ when compared with control PBS and double-deficient BMDC injection group (Fig. 8B-8D). This data suggested that T cell phenotypes required Sting expression in BMDC. Interestingly, only *Sting*-sufficient BMDC from *Fcgr2b*^-/-^, but not WT mice, induced the spontaneous germinal center B cell formation. Also, *Sting*-deficient BMDC from *Fcgr2b*^-/-^ did not increase germinal center B cell (Fig. 8D). Next, we examined the immunofluorescence staining to identify the immune complexes at the kidney of the recipient. The recipient of *Sting*-sufficient BMDC showed an increase of IgG deposition and CD45^+^ cell infiltration while *Sting*-deficient BMDC did not (Fig. 8E-8H). Nevertheless, the *Fcgr2b^-/-^* BMDC induced more immune complexes and CD45^+^ cells in the kidneys (Fig. 8F). The results suggested that the restoration of the STING-signaling pathway in dendritic cells is essential for lupus development in the *Fcgr2b^-/-^. Sting^gt/gt^* mice.

In summary, STING induced DC maturation to become professional APC and plasmacytoid DC via LYN kinase. The STING-activated DCs induced T cell proliferation and autoimmunity. Inhibition of STING pathway rescued lupus phenotypes.

## Discussion

A gain-of-function mutation in STING has been identified as a responsible gene for a subpopulation of SLE patients, and STING-dependent interferon-inducible genes correlated with disease activity ^38, 39^. However, there is no functional data of STING in a lupus mouse model that relevant to human SLE. The 129/B6.*Fcgr2b^-/-^* mice carrying Nba2 region expressed constitutively *Ifi202*. The CD19^+^ cells from B6.Nba2 shows the increase of *Ifi202* and the decreases of *Sting* expression ^40^. However, the overexpression of *Ifi202* can activate the Sting-dependent IFN-I response and the 129/B6.*Fcgr2b^-/-^* mice increase the expression of IFN-β ^41, 42^. Here, we detected the high expression of *Ifi202*, *Sting*, *Ifn-β*, and interferon-inducible genes from the spleen of these mice. Although STING functions as a negative regulator in the Mrl/*lpr* lupus mice ^22^, our data show that STING is required for the lupus development in the 129/B6.*Fcgr2b^-/-^* mice. STING may also play a crucial role in other lupus mouse models, which contained the Nba2 region.

The survival of the 129/B6.*Fcgr2b^-/-^* mice depend on autoantibody production and glomerulonephritis ^43, 44^. STING is required for the antibody production induced by cyclic-di-GMP in vitro ^45^. These data suggested that STING facilitated the autoantibody production, inflammatory cell infiltration, and glomerulonephritis in the 129/B6.*Fcgr2b^-/-^* mice. The expression of interferon-inducible genes associated with SLE disease activity ^46^. We detected the very high expression of IFN inducible genes in the kidneys of 129/B6.*Fcgr2b^-/-^* mice showed severe pathology. The absence of STING signaling in the *Fcgr2b^-/-^* mice partly decreased the expression of interferon-inducible genes in the kidney. This data suggested that other nucleic acid sensors may promote the type I interferon production or signaling in the *Fcgr2b^-/-^* mice as well, and STING-dependent lupus phenotypes do not mediate only through type-I interferon pathway.

STING expresses and functions differentially depended on the cell types. STING signals synergistically with B cell receptor signaling to promote antibody response ^47^. Our results showed that spontaneous germinal center B cells and MHC-II expression in the *Fcgr2b^-/-^* mice were *Sting*-dependent. However, plasma cell expansion was *Sting*-independent. This data suggested STING may contribute to the autoantibody production through memory B cells. STING also activates T cells, which induced type I IFN production and mediated cell death ^48^. Nevertheless, we found that the increase of T effector memory (Tem) in the *Fcgr2b^-/-^* mice was *Sting*-dependent. The expansion of Tem may directly mediate through the interaction with APC, not via Sting signaling in T cells.

STING agonist (DMXAA) treated mice show the increased expression of CD80, CD86, and MHC-II on DC, suggesting the mature phenotypes of DC as the APC ^49^. We observed the reduction of DC expansion in the *Fcgr2b^-/-^* mice, which depended on STING signaling. We confirmed that STING was required for DC maturation and cytokine production. These DC became professional APC and could promote T cell differentiation. The IFN-γ producing CD4^+^ cells in the spleen of the *Fcgr2b^-/-^* mice was reduced in the absence of STING. The *Sting*-expressing DC derived from WT and *Fcgr2b^-/-^* mice stimulated naïve T cells to proliferate, however, the ability of T cell to differentiate and produce IFN-γ did not depend on intrinsic *Sting* expression on T cells. Interestingly, only DC from the *Fcgr2b^-/-^* mice can increase the IFN-γ production in CD4^+^ T cells. This data suggested the DC from the *Fcgr2b^-/-^* mice have the intrinsic property that promotes the generation of IFN-γ producing CD4^+^ T cells.

The cGAS-STING signaling can activate human pDCs to produce IFN-I and knockdown of *Sting* using siRNA in CAL-1 cells can cause the reduction of IFN response ^50^. The proteomic data showed the upregulation of interferon-regulated protein after STING-activation with DMXAA, which implied that the culture environment should enrich with type-I IFN. STING activation led to phosphorylation of Ser357 of mouse Sting (homolog Ser358 in human Sting), and this site is phosphorylated by TBK1, which subsequently promoting type I IFN production ^51, 52^. The identification of pDC, a major producer of type-I IFN, after STING activation, uncovered the role of STING in the differentiation of pDC. These data revealed that STING was essential for the generation of pDCs. Besides, our study identified several STING-interacting proteins by mass spectrometry. Lyn kinase has been shown the role in the differentiation of pDC^37^. The recruitment of LYN to STING after DMXAA stimulation suggested STING mediated signaling through Lyn kinase. Also, the proteomics data of STING-activated BMDC showed a significant increase of phosphoinositide 3-kinase adapter protein 1 (Pik3ap1) and receptor of activated protein C kinase 1 (Rack1) (Supplemental Table 1). Pik3ap1 is an adaptor that signals to the phosphoinositide 3-kinase (PI3K) ^53^. LYN and RACK1 are co-immunoprecipitated in membrane complexes ^54^. RACK1 silencing effected on the phosphorylation of AKT ^55^. Our data suggest that the downstream of STING-LYN signaling mediated through the PI3K-AKT pathway. The inhibition of Lyn kinase during STING activation diminished DC maturation and pDC differentiation. The data suggested STING mediated differentiation of BMDC through the LYN signaling pathway.

The depletion of pDC ameliorates the autoimmune phenotypes in BXSB lupus-prone mice and B6.Nba2 mice ^56, 57^. Our data strongly suggested STING involving in DC function both DC maturation and pDC differentiation. Adoptive transfer of *Sting*-sufficient BMDCs can induce autoantibody production and immune complex deposition regardless of the *Fcgr2b* status. However, the absence of *Fcg2b* in the BMDC can accelerate the severity of autoimmune phenotypes, especially increased inflammatory cell infiltration in the kidney of the double-deficient recipient mice. These data suggested that STING-expression on BMDC is essential for the initiation of autoimmunity.

In conclusion, this study established the vital function of STING in the autoimmune *Fcgr2b^-/-^* lupus mouse model, thus providing a strong tool for future mechanistic and preclinical studies of STING in SLE. Inhibition of STING signaling represents a promising therapeutic target for SLE patients.

## Methods

### Mice

The *Fcgr2b*-deficient mice on the 129/C57BL/6 background were obtained from Dr. Bolland (NIH, Maryland, USA). *Sting*-deficient mice were provided by Paludan (Aarhus University, Aarhus, Denmark). Wild type mice were purchased from the National Laboratory Animal Center, Nakornpathom, Thailand. The *Fcgr2b*-deficient mice were crossed with *Sting*-deficient mice to generate the double-deficient mice and their littermate controls. The Animal Experimentation Ethics Committee of Chulalongkorn University Medical School approved the animal protocols.

### Measurement of autoantibody

Blood was collected from the mice at the age of 6-7 months. The levels of anti-dsDNA were measured by Enzyme-linked immunosorbent assay (ELISA). The anti-nuclear antibodies were detected by indirect immunofluorescence using HEp-2 cells (EUROIMMUN, Luebeck, Germany).

### Cytokine detection

Cytokine panels in serum including IL-1α, IL-1β, IL-6, IL-10, IL-12p70, IL-17A, IL-23, IL-27, MCP-1, IFN-β, IFN-γ, TNF-α, and GM-CSF were measured using LEGENDplex™ Mouse Inflammation Panel kit (Biolegend, San Diego, CA, USA). The beads were read on a flow cytometer using BDTM LSR-II (BD Biosciences, USA).

### Single-cell preparations

Splenocytes were isolated from all experimental groups and littermate wild-type mice. CD4^+^ T cells and naïve T cells were isolated using CD4 isolation Kit and naïve CD4^+^ T cell Isolation Kit (Miltenyi, Bergisch Gladbach, Germany).

### Flow cytometry

The splenocytes (1 x 10^6^ cells) were stained with flow antibody including anti-CD4 (GK1.5), CD8 (53-6.7), CD62L (MEL-14), CD44 (IM7), CD3ε (145-2C11), ICOS (C398.4A), CD11c (N418), B220 (RA3-6B2), CD11b (M1/70), I-Ab (AF6-120.1), PDCA-1 (129c1), CD80 (16-10A1), GL7 (GL7), CD138 (281-2), IFN-γ (XMG1.2) (Biolegend, San Diego, CA, USA) and Fixable Viability Dye eFluor® 780 (Thermo Fisher Scientific, MA USA). The flow cytometry was performed using BDTM LSR-II (BD Biosciences, USA) and analysis by FlowJo software (USA).

### Gene expression analysis

A total of RNA was extracted from the kidneys and spleen by Trizol reagent (Invitrogen, CA, USA) followed by RNA purified using the RNeasy mini kit and treated with DNase I (Qiagen, MD, USA). Then, one µg of total RNA was used as a template for cDNA synthesis using iScript RT Supermix (Biorad, California, USA). The expression of interest genes was assessed by quantitative real-time PCR. The primer sequences used in the real-time PCR show as follow; *Cxcl10*: CAGTGAGAATGAGGGCCATAGG and CGGATTCAGACATCTCTGCTCAT; *Mx1*: GATCCGACTTCACTTCCAGATGG and CATCTCAGTGGTAGTCCAACCC; *Irf7*: CCCAGACTGCCTGTGTAGACG and CCAGTCTCCAAACAGCACTCG; *Ifnb*: ATGAGTGGTGGTTGCAGGC and TGACCTTTCAAATGCAGTAGATTCA; *Ifng*: TTGCCAAGTTTGAGGTCAACAA and TGGTGGACCACTCGGATGA; *p202*: AGCCTCTCCTGGACCTAACA and GCAGTGAGTACCATCACTGTCA; *Sting*: TGCCGGACACTTGAGGAAAT and GTTTCCGTCTGTGGGTTCTTG; *Actb*; GGCTGTATTCCCCTCCATCG and CCAGTTGGTAACAATGCCATGT; *Isg15*: GAGCTAGAGCCTGCAGCAAT and TAAGACCGTCCTGGAGCACT; *Irf5*: TTTGAGATCTTCTTTTGCTTTGGA and GTACCACCTGTACAGTAATGAGCTTCTT. Microarray analysis was performed by RNA labeling and hybridization using the Agilent One-Color Microarray-Based Gene Expression Analysis protocol (Agilent Technology, V 6.5, 2010)

### Immunofluorescence and Histopathology

Frozen renal sections were fixed in acetone and blocked with 1% BSA in PBS. The sections were stained with FITC-conjugated goat anti-mouse IgG antibodies, PE-conjugated goat anti-mouse C45 antibodies (Abcam, Cambridge, MA, USA), and DAPI (4’,6-Diamidino-2-Phenylindole, Dihydrochloride) (Thermo Fisher Scientific, MA USA). The fluorescent signaling was visualized by ZEISS LSM 800 with Airyscan (Carl Zeiss, Germany). The deparaffinized kidney sections were fixed with formalin subsequently stained with H&E. The pathology grading from kidney sections was blinded analysis by the nephrologist.

### In vitro culture assays of BMDCs and T cells

Differentiation of bone marrow-derived dendritic cells (BMDC) was performed. T cells were labeled with CFSE (Supplementary method). Immature BMDCs were cultured and activated with DMXAA (5,6-Dimethylxanthenone-4-acetic acid or STING ligand) (Invivogen, San Diego, USA) for 24 hours at 37 °C. Activated BMDCs were plated with CFSE labeled T cells from lymph for 6 and 72 hours at 37 °C and 5% CO2 followed by intracellular staining for anti-IFN-γ.

### Colocalization of STING interacting proteins

Immature BMDCs from WT and *Fcgr2b^-/-^* were stimulated with DMXAA for 3 and 6 hours. Cells were fixed with 4% formalin (Sigma-Aldrich, Darmstadt, Germany) at room temperature for 15 minutes and stained with anti-LYN antibody (cat. 2732, Cell Signaling, MA, USA), anti-FYN antibody [FYN-01] (cat. 1881, Abcam, Cambridge, MA, USA), STING antibody conjugated with Zenon^TM^ Alexa fluor^TM^ 555 rabbit IgG labeling kit (Thermo Fisher Scientific, MA, USA), and DAPI (Thermo Fisher Scientific, MA USA). The fluorescent signaling was visualized by ZEISS LSM 800 with Airyscan (Carl Zeiss, Germany).

### Immunoprecipitation of STING-interacting proteins

BMDCs from WT, *Fcgr2b^-/-^* and *Sting^gt/gt^* were cultured, then stimulated with DMXAA for 3 and 6 hr. Cells were collected and lysed. The antibody was mixed with the magnetic beads by adding 10 µg of STING antibody (CUSB in-house antibody, clone: GTN-01; targeted N-terminal) with 400 µg of SureBeads™ Protein A magnetic beads (Biorad, California, USA) and incubated for 1 hour at room temperature. Then, Protein lysates were added and incubated with antibody-conjugated beads for overnight at 4 °C. The beads were washed 3 times, and samples were eluted. The eluted protein samples were separated by 10 % SDS-PAGE gel. The STING interacting proteins from co-IP were analyzed by in-gel digestion, followed by LC-MS/MS analysis.

### Western Blot Analysis

Splenocytes were lysed in 2 % SDS lysis buffer. Cell lysates containing protein (20 µg) were boiled in SDS sample buffer at 37 °C for 15 min before separation on a 10 % SDS-polyacrylamide gel. Proteins were transferred to nitrocellulose membranes and probed with STING antibody (clone: GTN-01; 1:2000) (CUSB-in house, BKK, TH) and LYN antibody (clone: LYN-01;1:1000) (Biolegend, San Diego, CA, USA). After incubation at 4 °C for overnight, the membrane was washed and probed with IRDye® 680RD Donkey anti-Rabbit IgG (H + L) and IRDye® 800CW Donkey anti-Mouse IgG (H + L) secondary antibody (1:10000) (LI-COR, Lincoln, Nebraska, USA) for 1 hour at room temperature. The membrane was determined the signals by ODYSSEY CLx (LI-COR, Lincoln, Nebraska, USA).

### Inhibition of LYN in Sting-activated BMDC

BMDCs were prepared as described above and stimulated with DMXAA. Cells were then incubated with the LYN inhibitors PP2 (Sigma-Aldrich, Darmstadt, Germany) for one hour before the addition of DMXAA and LPS for 24 hours. Total proteins from BMDC were analyzed by Western blotting using Phospho-LYN (Tyr507), LYN (C13F9), Phospho-AKT (Ser473), and CST-AKT antibody (Cell Signaling, MA, USA).

### Adoptive transfer

BMDCs from WT, *Fcgr2b^-/-^*, and *Fcgr2b^-/-^*.*Sting^gt/gt^* mice were transferred into the recipient WT or *Fcgr2b^-/-^*.*Sting^gt/gt^* mice (at the age of 4 months) every two weeks for 4 times.

### Statistical analysis

All statistical analyses employed the two-tailed Mann-Whitney test. Statistical analyses were performed using GraphPad Prism 4.0 (GraphPad Software, San Diego, CA).

## Online supplemental material

Fig. S1 shows the gene expression profiles of kidneys in the *Fcgr2b^-/-^* and *Fcgr2b^-/-^. Sting^gt/gt^* mice. Fig. S2 shows the immuno-phenotypes of the *Fcgr2b^-/-^*lupus mice and *Fcgr2b^-/-^. Sting^gt/gt^*.

Fig. S3 shows the phenotypes of STING-activated dendritic cells. Table S1 shows the list of STING-regulated proteins. Table S2 shows the list of STING-interacting proteins.

## Acknowledgments

We thank Silvia Bolland (NIH) for providing the 129/B6.*Fcgr2b^-/-^* mice.

This study was supported by the Thailand Research Fund (TRF) RSA5980023 to PP, International Network for Lupus Research (IRN59W0004) from TRF to PP, TP, and PW, the National Research Council of Thailand (NRCT) to Chulalongkorn University (2015**-**2018) to TP and PP, and Chulalongkorn Academic Advancement into the 2^nd^ Century (CUAASC) Project to TP.

The authors declare no competing financial interests.

## Author contributions

A. Thim-uam performed experiments, interpreted data, prepared the figures, and wrote the manuscript. A. Thim-uam performed most of the experiments. T. Prabakaran performed the in vitro functional assays with the inhibitors and prepared the figures. M. Tansakul, and T. Benjachat performed experimental assistance for flow cytometry. J. Makjaroen and P. Wongkongkathep performed proteomics experiments and analyses. N. Chantaravisoot provided experimental support for confocal microscopy. T. Saethang provided analysis assistance for microarray. A. Leelahavanichkul analyzed and scored histopathology of tissue sections. S.Paludan provided reagents and expertise related to Sting signaling, co**-**directed the study, and edited the manuscript. T.Pisitkun contributed reagents and expertise related to mass**-**spectrometry, co**-**directed the study, and edited the manuscript. P.Pisitkun performed experiments, interpreted data, directed the studies, and wrote the manuscript.

## Authorship note

A. Thim-uam currently works at Division of Biochemistry, School of the Medical Sciences University of Phayao, Phayao Thailand. M. Tansakul currently works at the Department of Clinical Sciences and Public Health, Faculty of Veterinary Science, Mahidol University, Nakhon Pathom, Thailand.

