## Supplementary Data for "Inhibition of Sting rescues lupus disease by the regulation of Lyn-mediated dendritic cell differentiation"

### Supplementary Materials:

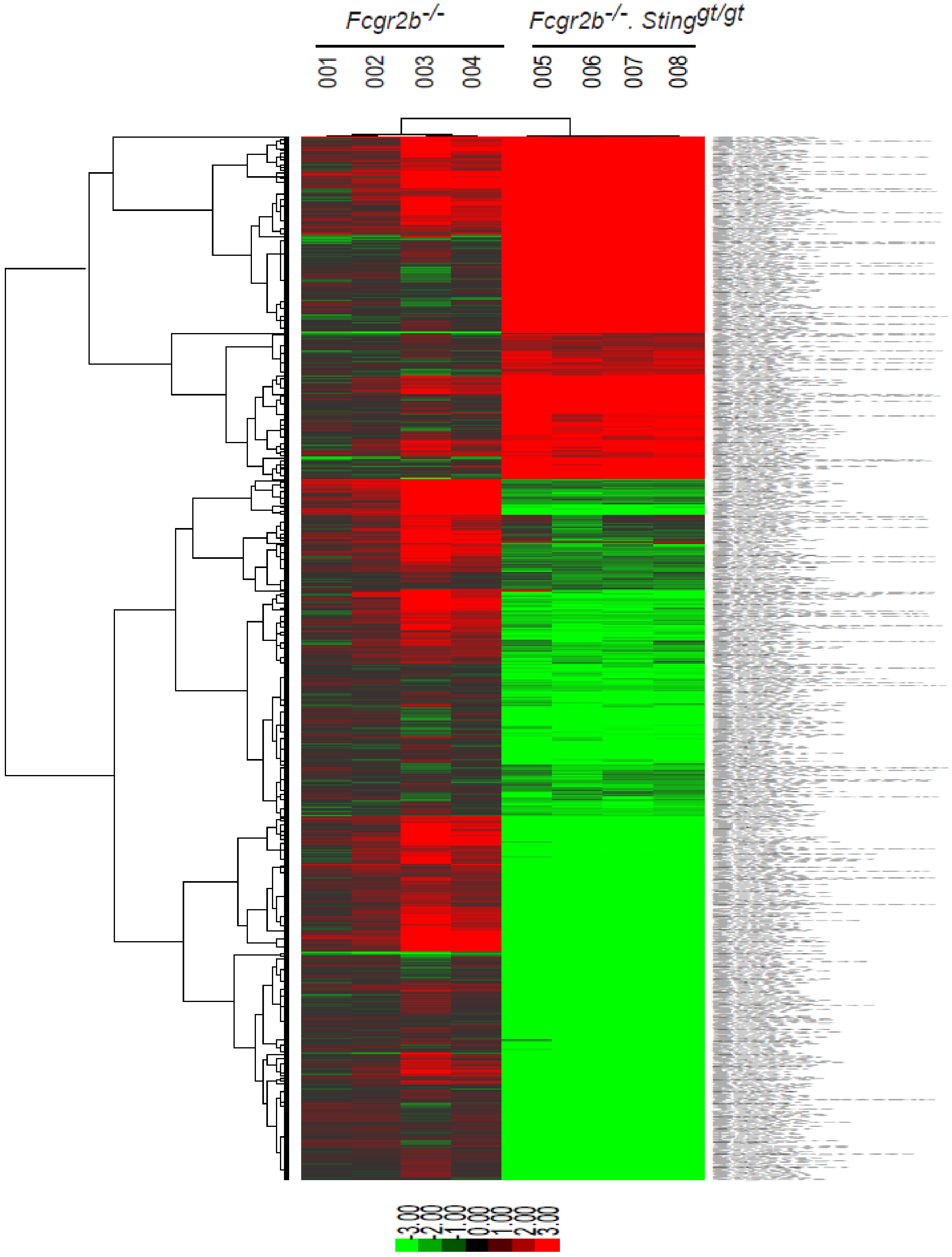


**Supplemental Figure 1. Gene expression profiles of kidneys in the *Fcgr2b*$^{-/-}$ and *Fcgr2b*$^{-/-}$. *Sting*$^{gt/gt}$ mice.** (A) A heat map of microarray data show the genes that significantly changed up to 2 fold compared between *Fcgr2b*$^{-/-}$ and *Fcgr2b*$^{-/-}$. *Sting*$^{gt/gt}$ mice (N=4 mice per group; $p<0.05$). Data show in log2 (sample/wild-type).

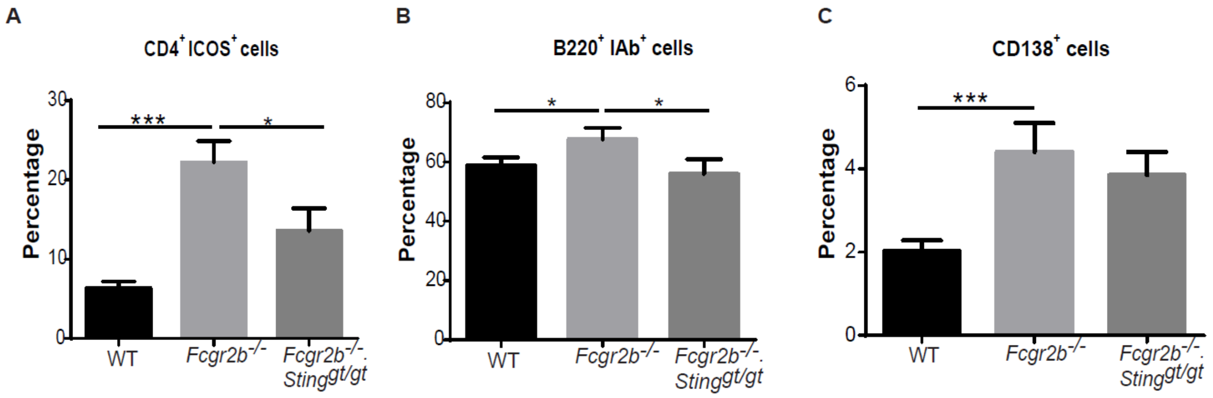


**Supplemental Figure 2. Sting signaling is essential for immuno-phenotypes of the *Fcgr2b*$^{-/-}$lupus mice**. (A-C) Flow cytometry analysis of splenocytes isolated from wild-type, *Fcgr2b*$^{-/-}$ and *Fcgr2b*$^{-/-}$. *Sting*$^{gt/gt}$ mice at the age of 6-7 months (N= 13-14 per group). Data shown in the percentage of (A) CD4$^{+}$ ICOS$^{+}$ cells, (B) B220$^{+}$ I-Ab$^{+}$ cells and (C) CD138$^{+}$ cells. Data show as mean ± SEM (*p < 0.05, **p<0.01 and ***p<0.001).

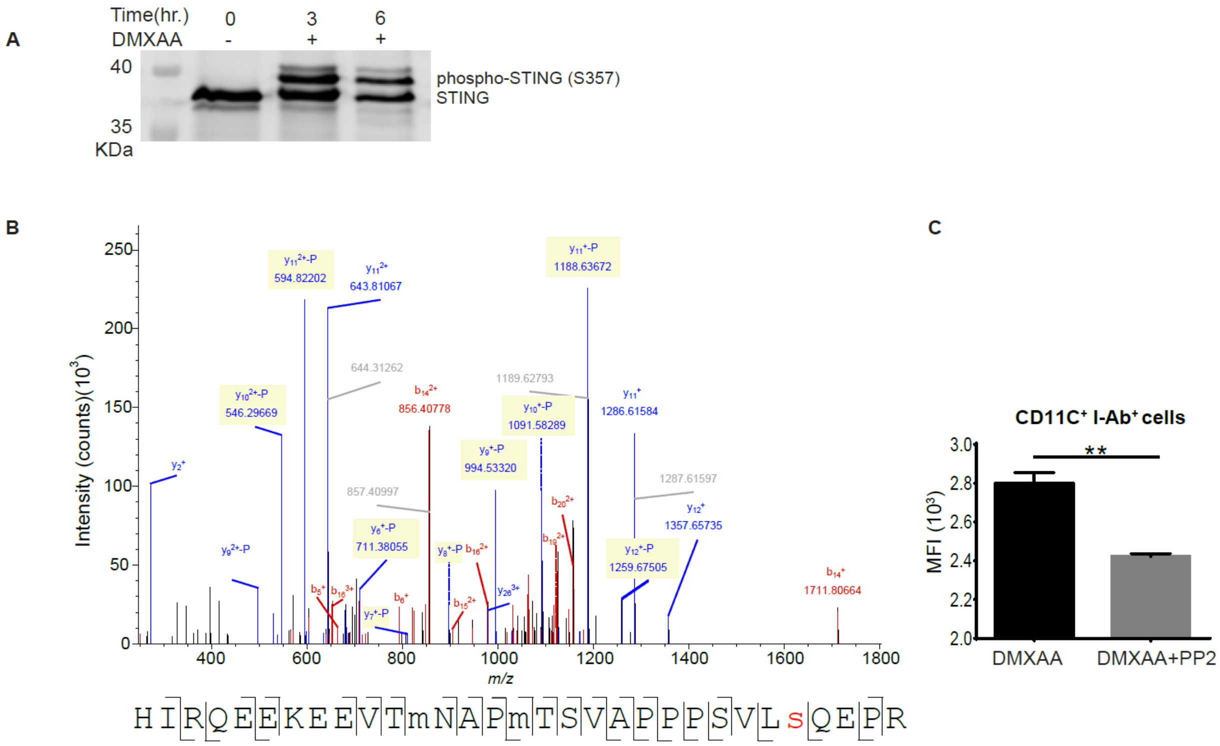


**Supplemental Figure 3. Phenotypes of Sting activated dendritic cells**. (A) Representative of western blot analysis from immunoprecipitation with Sting of *Fcgr2b*$^{-/-}$ mice (N= 4). The band was shown in STING protein of activated BMDC with DMXAA at 0, 3 and 6 hr. and phosphorylation of STING at Ser357. (B) Mass spectra of phosphorylation of STING at Ser357 of activated BMDC from *Fcgr2b*$^{-/-}$ mice after stimulated with DMXAA for 3 hour and followed by immunoprecipitation with STING. (C) Sting-activated BMDC were co-cultured with LYN inhibitor PP2 and analyzed by flow cytometry, which showed the mean fluorescence intensity (MFI) of IAb expressing DC (N = 3 mice per group).

### Supplemental Table 1. Lists of up and down of regulated proteins

| Accession No. | Gene ID | Description | *Fcgr2b*$^{-/-}$/*Fcgr2b*$^{-/-}$ .*Sting*$^{gt/gt}$(Log2) |
|---|---|---|---|
| Q8BGQ7 | Aars | Alanine--tRNA ligase, cytoplasmic | -0.217766032 |
| P48410 | Abcd1 | ATP-binding cassette sub-family D member 1 | -1.532028524 |
| Q99LE6 | Abcf2 | ATP-binding cassette sub-family F member 2 | -0.361706369 |
| Q99LR1 | Abhd12 | Monoacylglycerol lipase ABHD12 | -0.61955391 |
| Q8BWT1 | Acaa2 | 3-ketoacyl-CoA thiolase, mitochondrial | -0.881074395 |
| Q8JZN5 | Acad9 | Acyl-CoA dehydrogenase family member 9, mitochondrial | -0.325274114 |
| P45952 | Acadm | medium-chain specific acyl-CoA dehydrogenase, mitochondrial | -0.732129901 |
| P50544 | Acadvl | Very long-chain specific acyl-CoA dehydrogenase, mitochondrial | -0.987770943 |
| Q6ZQK5 | Acap2 | Arf-GAP with coiled-coil, ANK repeat and PH domain-containing protein 2 | 0.599553772 |
| Q8QZT1 | Acat1 | Acetyl-CoA acetyltransferase, mitochondrial | -0.456502772 |
| Q8CAY6 | Acat2 | Acetyl-CoA acetyltransferase, cytosolic | -0.673410757 |
| Q99KI0 | Aco2 | Aconitate hydratase, mitochondrial | -0.55368288 |
| P54987 | Acod1 | cis-aconitate decarboxylase | 2.995209528 |
| Q9R0H0 | Acox1 | Peroxisomal acyl-coenzyme A oxidase 1 | -1.197548502 |
| P41216 | Acsl1 | Long-chain-fatty-acid--CoA ligase 1 | 2.929355003 |
| Q9QUJ7 | Acsl4 | Long-chain-fatty-acid--CoA ligase 4 | 0.248147941 |
| P62737 | Acta2 | Actin, aortic smooth muscle | 0.423270416 |
| Q9Z0F8 | Adam17 | Disintegrin and metalloproteinase domain-containing protein 17 | -0.479046929 |
| P28650 | Adssl1 | Isoform 2 of Adenylosuccinate synthetase isozyme 1 | -0.428796169 |
| Q8K2K6 | Agfg1 | Arf-GAP domain and FG repeat-containing protein 1 | 0.713028959 |
| Q8C0I1 | Agps | Alkyldihydroxyacetonephosphate synthase, peroxisomal | -0.752037672 |
| Q68FL4 | Ahcyl2 | Putative adenosylhomocysteinase 3 | 0.501805729 |
| O08915 | Aip | AH receptor-interacting protein | -0.364411445 |
| Q9JII6 | Akr1a1 | alcohol dehydrogenase [NADP(+)] | 0.96922145 |
| Q571I9 | Aldh16a1 | Aldehyde dehydrogenase family 16 member A1 | -0.399737759 |
| Q62148 | Aldh1a2 | retinal dehydrogenase 2 | 1.743060723 |
| Q8R0Y6 | Aldh1l1 | Cytosolic 10-formyltetrahydrofolate dehydrogenase | 0.560522175 |
| P47738 | Aldh2 | Aldehyde dehydrogenase, mitochondrial | -0.79393871 |
| Q80VQ0 | Aldh3b1 | Aldehyde dehydrogenase family 3 member B1 | -0.521808257 |
| Q8CHT0 | Aldh4a1 | Delta-1-pyrroline-5-carboxylate dehydrogenase, mitochondrial | -0.689220634 |
| Q9JLJ2 | Aldh9a1 | 4-trimethylaminobutyraldehyde dehydrogenase | -0.295687187 |
| P39654 | Alox15 | arachidonate 15-lipoxygenase | 1.089325312 |
| O08583 | Alyref | THO complex subunit 4 | -0.465432076 |
| O08739 | Ampd3 | AMP deaminase 3 | 1.928165431 |
| B2RXR6 | Ankrd44 | Serine/threonine-protein phosphatase 6 regulatory ankyrin repeat subunit B | -0.828950371 |

| | | | |
|---|---|---|---|
| O35639 | Anxa3 | annexin A3 | -0.468124073 |
| P61967 | Ap1s1 | AP-1 complex subunit sigma-1A | 0.555153149 |
| O54774 | Ap3d1 | AP-3 complex subunit delta-1 | 0.357822744 |
| Q9JKC8 | Ap3m1 | ap-3 complex subunit mu-1 | 0.272159427 |
| Q9DCR2 | Ap3s1 | AP-3 complex subunit sigma-1 | 0.34446585 |
| P28352 | Apex1 | DNA-(apurinic or apyrimidinic site) lyase | -0.756191191 |
| Q78IK4 | Apool | MICOS complex subunit MIC27 | -0.102608847 |
| P08030 | Aprt | Adenine phosphoribosyltransferase | -0.218475924 |
| P61205 | Arf3 | ADP-ribosylation factor 3 | 0.269943477 |
| P61750 | Arf4 | ADP-ribosylation factor 4 | 0.766967391 |
| Q8BYW1 | Arhgap25 | Rho GTPase-activating protein 25 | -0.884585903 |
| Q3TBD2 | Arhgap45 | minor histocompatibility protein HA-1 | -0.536205649 |
| Q6P3A9 | Arl11 | ADP-ribosylation factor-like protein 11 | -0.502476419 |
| Q9JKW0 | Arl6ip1 | ADP-ribosylation factor-like protein 6-interacting protein 1 | 0.547476335 |
| Q8R5J9 | Arl6ip5 | PRA1 family protein 3 | -0.310735492 |
| Q9CQW2 | Arl8b | ADP-ribosylation factor-like protein 8B | -0.154095136 |
| Q9CVB6 | Arpc2 | Actin-related protein 2/3 complex subunit 2 | 0.224776872 |
| Q9WV54 | Asah1 | Acid ceramidase | -0.44823442 |
| E9PZJ8 | Ascc3 | Activating signal cointegrator 1 complex subunit 3 | 0.815581588 |
| O54984 | Asna1 | ATPase ASNA1 | -0.153053339 |
| P16460 | Ass1 | Argininosuccinate synthase | 1.262737415 |
| Q9CPX6 | Atg3 | ubiquitin-like-conjugating enzyme ATG3 | -0.643812087 |
| Q9D906 | Atg7 | Ubiquitin-like modifier-activating enzyme ATG7 | -0.295969125 |
| Q9CWJ9 | Atic | bifunctional purine biosynthesis protein purH | -0.365313119 |
| Q8VDN2 | Atp1a1 | Sodium/potassium-transporting ATPase subunit alpha-1 | -0.324374496 |
| Q6PIC6 | Atp1a3 | Sodium/potassium-transporting ATPase subunit alpha-3 | -1.075398517 |
| P97370 | Atp1b3 | sodium/potassium-transporting ATPase subunit beta-3 | -0.230978949 |
| Q64518 | Atp2a3 | Sarcoplasmic/endoplasmic reticulum calcium atpase 3 | -0.483845029 |
| Q06185 | Atp5i | ATP synthase subunit e, mitochondrial | -0.680497454 |
| P56135 | Atp5j2 | ATP synthase subunit f, mitochondrial | -0.437271018 |
| Q9DB20 | Atp5o | ATP synthase subunit O, mitochondrial | -0.412367771 |
| Q9Z1G4 | Atp6v0a1 | V-type proton ATPase 116 kDa subunit a isoform 1 | -0.706265175 |
| Q80SY3 | Atp6v0d2 | V-type proton ATPase subunit d 2 | -0.758413143 |
| P62814 | Atp6v1b2 | V-type proton ATPase subunit B, brain isoform | -0.274012334 |
| Q9JLZ3 | Auh | methylglutaconyl-CoA hydratase, mitochondrial | -0.64766609 |
| Q09200 | B4galnt1 | Beta-1,4 N-acetylgalactosaminyltransferase 1 | -1.645842095 |
| Q60739 | Bag1 | BAG family molecular chaperone regulator 1 | 0.941670817 |
| Q91XV3 | Basp1 | Brain acid soluble protein 1 | 0.898069323 |
| Q9Z277 | Baz1b | Tyrosine-protein kinase BAZ1B | -0.594705857 |
| O35855 | Bcat2 | Branched-chain-amino-acid aminotransferase, mitochondrial | -0.543188586 |

| | | | |
|---|---|---|---|
| Q8K019 | Bclaf1 | Bcl-2-associated transcription factor 1 | 0.174869878 |
| Q80XN0 | Bdh1 | D-beta-hydroxybutyrate dehydrogenase, mitochondrial | -1.606425093 |
| Q8R016 | Blmh | bleomycin hydrolase | -0.267550595 |
| Q9CY64 | Blvra | Biliverdin reductase A | -0.447422089 |
| Q9Z0S1 | Bpnt1 | 3'(2'),5'-bisphosphate nucleotidase 1 | 0.393595374 |
| Q7JJ13 | Brd2 | Bromodomain-containing protein 2 | 0.811632628 |
| Q8K2Q7 | Brox | BRO1 domain-containing protein BROX | 0.990089677 |
| P18572 | Bsg | Basigin | 1.162129002 |
| Q8R2Q8 | Bst2 | bone marrow stromal antigen 2 | 4.966481379 |
| P01027 | C3 | Complement C3 | 0.699329386 |
| Q64444 | Ca4 | Carbonic anhydrase 4 | 0.726283017 |
| Q9CR86 | Carhsp1 | Calcium-regulated heat stable protein 1 | 0.459788301 |
| P29452 | Casp1 | caspase-1 | 0.868079298 |
| P97864 | Casp7 | Caspase-7 | 1.012874895 |
| O89110 | Casp8 | Caspase-8 | 0.543221573 |
| P24270 | Cat | catalase | -0.541070258 |
| P23198 | Cbx3 | chromobox protein homolog 3 | -0.249126147 |
| Q9JIG7 | Ccdc22 | Coiled-coil domain-containing protein 22 | -0.473418781 |
| P47774 | Ccr7 | C-C chemokine receptor type 7 | 1.423253149 |
| P80318 | Cct3 | T-complex protein 1 subunit gamma | -0.216208222 |
| P11609 | Cd1d1 | Antigen-presenting glycoprotein CD1d1 | 1.308959057 |
| Q9ES57 | Cd200r1 | Cell surface glycoprotein CD200 receptor 1 | -1.376107182 |
| Q9EP73 | Cd274 | Programmed cell death 1 ligand 1 | 2.759549661 |
| Q9CWK3 | Cd2bp2 | CD2 antigen cytoplasmic tail-binding protein 2 | 0.364137601 |
| Q08857 | Cd36 | Platelet glycoprotein 4 | -1.087856028 |
| P56528 | Cd38 | ADP-ribosyl cyclase/cyclic ADP-ribose hydrolase 1 | 1.917522811 |
| P27512 | Cd40 | Tumor necrosis factor receptor superfamily member 5 | 3.61718323 |
| P41731 | Cd63 | CD63 antigen | -0.88845899 |
| P31996 | Cd68 | Macrosialin | -0.618281482 |
| P04441 | Cd74 | H-2 class II histocompatibility antigen gamma chain | -0.901080264 |
| P04441 | Cd74 | Isoform Short of H-2 class II histocompatibility antigen gamma chain | -0.901080264 |
| P42082 | Cd86 | T-lymphocyte activation antigen CD86 | 3.842512849 |
| Q9Z0M6 | Cd97 | CD97 antigen | -1.902973329 |
| Q61081 | Cdc37 | Hsp90 co-chaperone Cdc37 | 0.526181397 |
| Q8JZM7 | Cdc73 | Parafibromin | -0.350073064 |
| P24788 | Cdk11b | Cyclin-dependent kinase 11B | 0.36409994 |
| P97377 | Cdk2 | cyclin-dependent kinase 2 | -1.085690093 |
| P28033 | Cebpb | CCAAT/enhancer-binding protein beta | 1.365460565 |
| Q9Z0H4 | Celf2 | Isoform 9 of CUGBP Elav-like family member 2 | -0.454500617 |
| P11680 | Cfp | Properdin | -1.461066675 |
| Q9D1L0 | Chchd2 | Coiled-coil-helix-coiled-coil-helix domain-containing protein 2 | -0.94187026 |

| | | | |
|---|---|---|---|
| Q9CRB9 | Chchd3 | MICOS complex subunit MIC19 | -0.483197538 |
| Q8WTY4 | Ciapin1 | Anamorsin | 0.629707034 |
| Q8BRT1 | Clasp2 | CLIP-associating protein 2 | 1.104060091 |
| Q8BRT1 | Clasp2 | Isoform 2 of CLIP-associating protein 2 | 1.104060091 |
| Q9QZ15 | Clec4a | C-type lectin domain family 4 member A | -1.489388017 |
| Q6QLQ4 | Clec7a | C-type lectin domain family 7 member A | -1.663326202 |
| Q9QYB1 | Clic4 | Chloride intracellular channel protein 4 | 1.831008014 |
| Q8VBZ3 | Clptm1 | cleft lip and palate transmembrane protein 1 homolog | 0.23712684 |
| Q5SW19 | Cluh | Clustered mitochondria protein homolog | -1.007282341 |
| Q3U5Q7 | Cmpk2 | UMP-CMP kinase 2, mitochondrial | 3.628148861 |
| P53996 | Cnbp | Isoform 3 of Cellular nucleic acid-binding protein | -0.480058233 |
| P53996 | Cnbp | Cellular nucleic acid-binding protein | -0.480058233 |
| Q9D1A2 | Cndp2 | cytosolic non-specific dipeptidase | 0.353940959 |
| Q8BQ47 | Cnpy4 | Protein canopy homolog 4 | -0.40511594 |
| Q8K297 | Colgalt1 | Procollagen galactosyltransferase 1 | -0.348956756 |
| O88587 | Comt | Catechol O-methyltransferase | -0.734098424 |
| Q8CIE6 | Copa | coatomer subunit alpha | 0.087583893 |
| Q9JIF7 | Copb1 | Coatomer subunit beta | 0.123221328 |
| P61202 | Cops2 | Isoform 2 of COP9 signalosome complex subunit 2 | -0.721886983 |
| Q8C0P5 | Coro2a | Coronin-2A | -0.51921812 |
| Q9CQI6 | Cotl1 | coactosin-like protein | -0.60281177 |
| Q8BH51 | Cox14 | Cytochrome c oxidase assembly protein COX14 | 0.589568955 |
| Q9D7J4 | Cox20 | Cytochrome c oxidase protein 20 homolog | -0.874035806 |
| P19783 | Cox4i1 | Cytochrome c oxidase subunit 4 isoform 1, mitochondrial | -0.462660325 |
| P12787 | Cox5a | Cytochrome c oxidase subunit 5A, mitochondrial | -0.539015557 |
| P43024 | Cox6a1 | Cytochrome c oxidase subunit 6A1, mitochondrial | -1.596608351 |
| P56391 | Cox6b1 | Cytochrome c oxidase subunit 6B1 | -0.645262368 |
| P48771 | Cox7a2 | Cytochrome c oxidase subunit 7A2, mitochondrial | -0.760161865 |
| Q7TN98 | Cpeb4 | Cytoplasmic polyadenylation element-binding protein 4 | 2.678143292 |
| Q6NVF9 | Cpsf6 | Cleavage and polyadenylation specificity factor subunit 6 | 0.412168073 |
| O88668 | Creg1 | Protein CREG1 | -0.740480794 |
| P97315 | Csrp1 | Cysteine and glycine-rich protein 1 | 0.630873838 |
| O89098 | Cst7 | Cystatin-F | 1.426639832 |
| Q62426 | Cstb | Cystatin-B | 0.473935379 |
| Q61164 | Ctcf | Transcriptional repressor CTCF | -0.447563827 |
| P30999 | Ctnnd1 | Catenin delta-1 | 0.148385233 |
| P16675 | Ctsa | lysosomal protective protein | -0.877147749 |
| P49935 | Ctsh | Pro-cathepsin H | -1.35483901 |
| O70370 | Ctss | cathepsin S | -0.960048347 |
| Q9WTX6 | Cul1 | Cullin-1 | 0.887188394 |

| | | | |
|---|---|---|---|
| Q9D0M3 | Cyc1 | Cytochrome c1, heme protein, mitochondrial | -0.52725127 |
| Q80YW0 | Cyth4 | Cytohesin-4 | -0.936134618 |
| Q91WC9 | Daglb | Sn1-specific diacylglycerol lipase beta | -3.013510721 |
| Q9ER88 | Dap3 | 28S ribosomal protein S29, mitochondrial | -1.117515654 |
| P31786 | Dbi | acyl-CoA-binding protein | -0.478051358 |
| P43346 | Dck | deoxycytidine kinase | 2.523853652 |
| Q91YD3 | Dcp1a | mRNA-decapping enzyme 1A | 0.756554201 |
| Q8CBY8 | Dctn4 | Dynactin subunit 4 | 0.281329745 |
| Q80YA3 | Ddhd1 | Phospholipase DDHD1 | 0.406839382 |
| Q91VR5 | Ddx1 | ATP-dependent RNA helicase DDX1 | -0.148356236 |
| Q9JIK5 | Ddx21 | Nucleolar RNA helicase 2 | -0.550861152 |
| Q62167 | Ddx3x | ATP-dependent RNA helicase DDX3X | -0.369473886 |
| Q61656 | Ddx5 | probable ATP-dependent RNA helicase DDX5 | -0.489332367 |
| Q6Q899 | Ddx58 | Probable ATP-dependent RNA helicase DDX58 | 2.585247101 |
| Q9CQ62 | Decr1 | 2,4-dienoyl-CoA reductase, mitochondrial | -0.595262516 |
| O09005 | Degs1 | Sphingolipid delta(4)-desaturase DES1 | -1.20642232 |
| P00375 | Dhfr | dihydrofolate reductase | -0.46593848 |
| Q99J87 | Dhx58 | probable ATP-dependent RNA helicase DHX58 | 3.068421711 |
| O70133 | Dhx9 | Isoform 2 of ATP-dependent RNA helicase A | -0.402451588 |
| O08808 | Diaph1 | Protein diaphanous homolog 1 | 0.323589337 |
| Q9ESX5 | Dkc1 | H/ACA ribonucleoprotein complex subunit 4 | -0.320093611 |
| B1AZP2 | Dlgap4 | Disks large-associated protein 4 | 1.270065741 |
| Q9D2G2 | Dlst | Dihydrolipoyllysine-residue succinyltransferase component of 2-oxoglutarate dehydrogenase complex, mitochondrial | -0.502714216 |
| P63037 | Dnaja1 | DnaJ homolog subfamily A member 1 | 0.866039391 |
| Q99M87 | Dnaja3 | DnaJ homolog subfamily A member 3, mitochondrial | -0.762538478 |
| Q91YW3 | Dnajc3 | DnaJ homolog subfamily C member 3 | -0.364199549 |
| Q9QYI3 | Dnajc7 | DnaJ homolog subfamily C member 7 | 0.190932198 |
| P56542 | Dnase2 | Deoxyribonuclease-2-alpha | -1.016178752 |
| P13864 | Dnmt1 | DNA (cytosine-5)-methyltransferase 1 | -0.437379785 |
| O88508 | Dnmt3a | DNA (cytosine-5)-methyltransferase 3A | -1.146451309 |
| Q8BZN6 | Dock10 | Isoform 3 of Dedicator of cytokinesis protein 10 | 0.884103185 |
| Q8C3J5 | Dock2 | Dedicator of cytokinesis protein 2 | -0.277522568 |
| Q8C147 | Dock8 | Dedicator of cytokinesis protein 8 | -0.300890858 |
| P97465 | Dok1 | docking protein 1 | 0.946400261 |
| O70469 | Dok2 | Docking protein 2 | -0.783200003 |
| Q99KK7 | Dpp3 | dipeptidyl peptidase 3 | -0.19212012 |
| Q9ET22 | Dpp7 | Dipeptidyl peptidase 2 | -0.846316437 |
| P32233 | Drg1 | developmentally-regulated GTP-binding protein 1 | 0.570924409 |
| Q3UIR3 | Dtx3l | E3 ubiquitin-protein ligase DTX3L | 2.175866085 |
| Q9JHU4 | Dync1h1 | Cytoplasmic dynein 1 heavy chain 1 | 0.293939022 |

| | | | |
|---|---|---|---|
| P62627 | Dynlrb1 | Dynein light chain roadblock-type 1 | 0.21434794 |
| P51807 | Dynlt1 | Dynein light chain Tctex-type 1 | 0.3492403 |
| O35459 | Ech1 | Delta(3,5)-Delta(2,4)-dienoyl-CoA isomerase, mitochondrial | -0.564174762 |
| Q8BH95 | Echs1 | Enoyl-CoA hydratase, mitochondrial | -0.522470078 |
| Q9WUR2 | Eci2 | Enoyl-CoA delta isomerase 2, mitochondrial | -1.061080602 |
| Q3UJB9 | Edc4 | Enhancer of mRNA-decapping protein 4 | -0.346058963 |
| O08810 | Eftud2 | 116 kDa U5 small nuclear ribonucleoprotein component | -0.310111415 |
| Q9WVK4 | Ehd1 | EH domain-containing protein 1 | 0.638747707 |
| Q8BJW6 | Eif2a | Eukaryotic translation initiation factor 2A | -0.347499944 |
| Q03963 | Eif2ak2 | interferon-induced, double-stranded RNA-activated protein kinase | 2.350926637 |
| Q99LD9 | Eif2b2 | Translation initiation factor eIF-2B subunit beta | 0.563147005 |
| Q9Z0N1 | Eif2s3x | Eukaryotic translation initiation factor 2 subunit 3, X-linked | 0.490987239 |
| O70194 | Eif3d | Eukaryotic translation initiation factor 3 subunit D | -0.254856295 |
| Q91WK2 | Eif3h | Eukaryotic translation initiation factor 3 subunit H | 0.395411095 |
| Q99JX4 | Eif3m | Eukaryotic translation initiation factor 3 subunit M | -1.020752337 |
| P10630 | Eif4a2 | Isoform 2 of Eukaryotic initiation factor 4A-II | 0.981899047 |
| Q91VC3 | Eif4a3 | Eukaryotic initiation factor 4A-III | -0.233549795 |
| Q62448 | Eif4g2 | Eukaryotic translation initiation factor 4 gamma 2 | 0.255131753 |
| Q9WUK2 | Eif4h | Eukaryotic translation initiation factor 4H | -0.30601965 |
| Q05D44 | Eif5b | Eukaryotic translation initiation factor 5B | 0.138114373 |
| O55135 | Eif6 | eukaryotic translation initiation factor 6 | 0.465848409 |
| Q8BHL5 | Elmo2 | Engulfment and cell motility protein 2 | -0.304723269 |
| P83940 | Eloc | Transcription elongation factor B polypeptide 1 | 0.158093974 |
| P21995 | Emb | Embigin | -0.671094957 |
| O35130 | Emg1 | Ribosomal RNA small subunit methyltransferase Nep1 | -0.987484118 |
| Q9JIX0 | Eny2 | Transcription and mRNA export factor ENY2 | -0.889405431 |
| Q8CGC7 | Eprs | Bifunctional glutamate/proline--tRNA ligase | 0.060870724 |
| P42567 | Eps15 | epidermal growth factor receptor substrate 15 | 0.548087647 |
| Q60902 | Eps15l1 | Epidermal growth factor receptor substrate 15-like 1 | 0.324403927 |
| Q3UVK0 | Ermp1 | Endoplasmic reticulum metallopeptidase 1 | 0.750541074 |
| Q9R0P3 | Esd | S-formylglutathione hydrolase | 0.809054086 |
| Q3U7R1 | Esyt1 | Extended synaptotagmin-1 | -0.50653897 |
| Q3TZZ7 | Esyt2 | Extended synaptotagmin-2 | -0.588347009 |
| Q921G7 | Etfdh | Electron transfer flavoprotein-ubiquinone oxidoreductase, mitochondrial | -1.016324838 |
| Q9DCM0 | Ethe1 | Persulfide dioxygenase ETHE1, mitochondrial | -0.703857428 |
| Q8VD58 | Evi2b | Protein EVI2B | -1.550419659 |
| Q8R3S6 | Exoc1 | exocyst complex component 1 | 0.529728685 |
| Q8BTW3 | Exosc6 | Exosome complex component MTR3 | -0.778610367 |
| Q9D753 | Exosc8 | Exosome complex component RRP43 | -0.773680555 |

| | | | |
|---|---|---|---|
| Q05816 | Fabp5 | Fatty acid-binding protein, epidermal | -0.214082441 |
| Q922J9 | Far1 | Isoform 4 of Fatty acyl-CoA reductase 1 | -1.569782283 |
| P19096 | Fasn | Fatty acid synthase | -0.447722091 |
| P35550 | Fbl | rRNA 2'-O-methyltransferase fibrillarin | -0.422648948 |
| P20491 | Fcer1g | High affinity immunoglobulin epsilon receptor subunit gamma | 0.90383595 |
| P08508 | Fcgr3 | Low affinity immunoglobulin gamma Fc region receptor III | -0.582677239 |
| P39749 | Fen1 | Flap endonuclease 1 | -0.92778771 |
| P97807 | Fh | fumarate hydratase, mitochondrial | -0.389669024 |
| P45878 | Fkbp2 | Peptidyl-prolyl cis-trans isomerase FKBP2 | -0.286717251 |
| Q62446 | Fkbp3 | peptidyl-prolyl cis-trans isomerase FKBP3 | -0.293707791 |
| Q8BTM8 | Flna | Filamin-A | -0.296438167 |
| Q80X90 | Flnb | Filamin-B | 1.757871636 |
| O08917 | Flot1 | Flotillin-1 | -1.176671894 |
| Q60634 | Flot2 | Flotillin-2 | -0.586462896 |
| A2APV2 | Fmnl2 | Isoform 3 of Formin-like protein 2 | 1.797624775 |
| P35922 | Fmr1 | Fragile X mental retardation protein 1 homolog | 0.489814745 |
| P11276 | Fn1 | fibronectin | 0.721873745 |
| Q8BX90 | Fndc3a | Fibronectin type-III domain-containing protein 3a | 1.338537846 |
| Q8K385 | FRRS1 | Ferric-chelate reductase 1 | -1.166363966 |
| Q91WJ8 | Fubp1 | Isoform 2 of Far upstream element-binding protein 1 | -0.258145539 |
| P97855 | G3bp1 | Ras GTPase-activating protein-binding protein 1 | -0.395081947 |
| P97379 | G3bp2 | Ras GTPase-activating protein-binding protein 2 | 0.383444047 |
| Q8K157 | Galm | aldose 1-epimerase | 0.299501857 |
| Q8BHN3 | Ganab | Isoform 2 of Neutral alpha-glucosidase AB | -0.496163105 |
| P16858 | Gapdh | glyceraldehyde-3-phosphate dehydrogenase | 0.094330465 |
| Q9CY66 | Gar1 | H/ACA ribonucleoprotein complex subunit 1 | -1.58702842 |
| Q9CZD3 | Gars | Glycine--tRNA ligase | 0.117047826 |
| Q64737 | Gart | trifunctional purine biosynthetic protein adenosine-3 | -0.504694802 |
| Q9Z0E6 | Gbp2 | Guanylate-binding protein 1 | 2.91571745 |
| Q61107 | Gbp4 | Guanylate-binding protein 4 | 4.460303796 |
| Q8CFB4 | Gbp5 | Guanylate-binding protein 5 | 5.073691882 |
| Q8R2Q4 | Gfm2 | Ribosome-releasing factor 2, mitochondrial | -0.956491178 |
| P47856 | Gfpt1 | glutamine--fructose-6-phosphate aminotransferase [isomerizing] 1 | 0.536029045 |
| Q9D7X8 | Ggct | gamma-glutamylcyclotransferase | 1.108596791 |
| Q9Z0L8 | Ggh | Gamma-glutamyl hydrolase | -0.739896736 |
| Q9Z0G0 | Gipc1 | PDZ domain-containing protein GIPC1 | -0.159012245 |
| Q9JLQ2 | Git2 | ARF GTPase-activating protein GIT2 | -0.35318862 |
| P23780 | Glb1 | Beta-galactosidase | -0.519964415 |
| Q61543 | Glg1 | Golgi apparatus protein 1 | -2.192293975 |
| Q60648 | Gm2a | Ganglioside GM2 activator | -0.515992005 |
| Q8BTZ7 | Gmppb | Mannose-1-phosphate guanyltransferase beta | 0.77651352 |

| | | | |
|---|---|---|---|
| P21279 | Gnaq | Guanine nucleotide-binding protein G(Q) subunit alpha | 0.437030744 |
| P63213 | Gng2 | Guanine nucleotide-binding protein G(I)/G(S)/G(O) subunit gamma-2 | -0.893952234 |
| Q921M4 | Golga2 | Golgin subfamily A member 2 | -0.306864366 |
| Q9QYE6 | Golga5 | Golgin subfamily A member 5 | 0.91327472 |
| Q99JX3 | Gorasp2 | Golgi reassembly-stacking protein 2 | -0.860138401 |
| P05202 | Got2 | Aspartate aminotransferase, mitochondrial | -0.363363977 |
| Q3U1Z5 | Gpsm3 | G-protein-signaling modulator 3 | -0.506106526 |
| P11352 | Gpx1 | Glutathione peroxidase 1 | -0.501438599 |
| Q99MK8 | Grk2 | Beta-adrenergic receptor kinase 1 | -1.084343813 |
| Q9D8T2 | Gsdmdc1 | Gasdermin-D | 0.691515669 |
| P10649 | Gstm1 | Glutathione S-transferase Mu 1 | -0.539407548 |
| O08582 | Gtpbp1 | GTP-binding protein 1 | -0.468524773 |
| P12265 | Gusb | beta-glucuronidase | -0.985095381 |
| Q80SU7 | Gvin1 | Interferon-induced very large GTPase 1 | 1.183837801 |
| Q9R062 | Gyg1 | Glycogenin-1 | 0.25707049 |
| Q9Z1E4 | Gys1 | glycogen [starch] synthase, muscle | 0.915551766 |
| P27661 | H2afx | Histone H2AX | -0.494792767 |
| P0C0S6 | H2afz | Histone H2A.Z | -0.604054662 |
| P01899 | H2-D1 | H-2 class I histocompatibility antigen, D-B alpha chain | 1.317256772 |
| P28078 | H2-DMa | Class II histocompatibility antigen, M alpha chain | -1.035640844 |
| P35737 | H2-DMb1 | Class II histocompatibility antigen, M beta 1 chain | -0.971781929 |
| P01901 | H2-K1 | H-2 class I histocompatibility antigen, K-B alpha chain | 1.052795605 |
| P06339 | H2-T23 | H-2 class I histocompatibility antigen, D-37 alpha chain | 0.9640568 |
| Q61425 | Hadh | Hydroxyacyl-coenzyme A dehydrogenase, mitochondrial | -0.811768415 |
| Q8BMS1 | Hadha | Trifunctional enzyme subunit alpha, mitochondrial | -0.501294703 |
| Q99JY0 | Hadhb | Trifunctional enzyme subunit beta, mitochondrial | -0.598582129 |
| Q99KB8 | Hagh | Hydroxyacylglutathione hydrolase, mitochondrial | -0.410894802 |
| P08103 | Hck | Tyrosine-protein kinase HCK | 0.434994598 |
| P49710 | Hcls1 | Hematopoietic lineage cell-specific protein | 0.347054528 |
| P70288 | Hdac2 | Histone deacetylase 2 | -0.499942841 |
| P51859 | Hdgf | hepatoma-derived growth factor | -0.263469895 |
| Q9R257 | Hebp1 | Heme-binding protein 1 | -0.875177244 |
| E9QAM5 | Helz2 | Helicase with zinc finger domain 2 | 2.216874888 |
| P20060 | Hexb | Beta-hexosaminidase subunit beta | -0.529939151 |
| Q3UDW8 | Hgsnat | heparan-alpha-glucosaminide N-acetyltransferase | -1.649386892 |
| Q9D0S9 | Hint2 | Histidine triad nucleotide-binding protein 2, mitochondrial | -0.595416352 |
| P43276 | Hist1h1b | Histone H1.5 | -0.416747479 |
| P43277 | Hist1h1d | Histone H1.3 | -0.399634767 |
| Q8CGP2 | Hist1h2bp | Isoform 2 of Histone H2B type 1-P | -0.460125705 |

| | | | |
|---|---|---|---|
| P62806 | Hist1h4a | histone H4 | -0.291751009 |
| P17710 | Hk1 | Hexokinase-1 | -0.236536725 |
| O08528 | Hk2 | Hexokinase-2 | 0.253952363 |
| Q3TRM8 | Hk3 | Hexokinase-3 | 0.729882645 |
| P17095 | Hmga1 | High mobility group protein HMG-I/HMG-Y | 1.120719228 |
| P38060 | Hmgcl | Hydroxymethylglutaryl-CoA lyase, mitochondrial | -0.692297943 |
| P14901 | Hmox1 | heme oxygenase 1 | 1.486475098 |
| Q9CX86 | Hnrnpa0 | Heterogeneous nuclear ribonucleoprotein A0 | -0.511089604 |
| Q9Z2X1 | Hnrnpf | Heterogeneous nuclear ribonucleoprotein F | -0.319800887 |
| Q9D0E1 | Hnrnpm | Heterogeneous nuclear ribonucleoprotein M | -0.078946288 |
| Q8VEK3 | Hnrnpu | Heterogeneous nuclear ribonucleoprotein U | -0.244414712 |
| Q3TEA8 | Hp1bp3 | Heterochromatin protein 1-binding protein 3 | -0.154788317 |
| Q6YGZ1 | Hpse | Heparanase | 0.324505715 |
| Q3TC93 | Hs1bp3 | HCLS1-binding protein 3 | -0.59953724 |
| O08756 | Hsd17b10 | 3-hydroxyacyl-CoA dehydrogenase type-2 | -0.417650472 |
| P51660 | Hsd17b4 | peroxisomal multifunctional enzyme type 2 | -0.632942451 |
| Q2TPA8 | Hsdl2 | Hydroxysteroid dehydrogenase-like protein 2 | -1.165724577 |
| P11499 | Hsp90ab1 | Heat shock protein HSP 90-beta | 0.351807004 |
| P17879 | Hspa1b | Heat shock 70 kDa protein 1B | 3.180820036 |
| Q7TMY8 | Huwe1 | E3 ubiquitin-protein ligase HUWE1 | -0.264505776 |
| P13597 | Icam1 | Intercellular adhesion molecule 1 | 1.921465693 |
| O88844 | Idh1 | Isocitrate dehydrogenase [NADP] cytoplasmic | -0.72392385 |
| P54071 | Idh2 | Isocitrate dehydrogenase [NADP], mitochondrial | -0.407267648 |
| Q9D6R2 | Idh3a | Isocitrate dehydrogenase [NAD] subunit alpha, mitochondrial | -0.672717546 |
| P0DOV2 | Ifi204 | interferon-activable protein 204 | 3.97429376 |
| Q9D8C4 | Ifi35 | Interferon-induced 35 kDa protein homolog | 1.475099066 |
| Q8R5F7 | Ifih1 | Interferon-induced helicase C domain-containing protein 1 | 3.299138366 |
| Q64282 | Ifit1 | interferon-induced protein with tetratricopeptide repeats 1 | 5.774506745 |
| Q64112 | Ifit2 | Interferon-induced protein with tetratricopeptide repeats 2 | 5.723988926 |
| Q64345 | Ifit3 | Interferon-induced protein with tetratricopeptide repeats 3 | 7.561106578 |
| Q9CQW9 | Ifitm3 | Interferon-induced transmembrane protein 3 | 2.678835336 |
| Q61249 | Igbp1 | Immunoglobulin-binding protein 1 | 0.703044083 |
| Q07113 | Igf2r | Cation-independent mannose-6-phosphate receptor | -0.773120212 |
| P25085 | Il1rn | Interleukin-1 receptor antagonist protein | 2.554381916 |
| O55222 | Ilk | Integrin-linked protein kinase | -0.59745988 |
| Q9ES52 | Inpp5d | Phosphatidylinositol 3,4,5-trisphosphate 5-phosphatase 1 | -0.775730444 |
| Q9EPL8 | Ipo7 | Importin-7 | -0.478093461 |
| Q64287 | Irf4 | Interferon regulatory factor 4 | 1.372738263 |
| P56477 | Irf5 | interferon regulatory factor 5 | 0.52120183 |
| Q60766 | Irgm1 | Immunity-related GTPase family M protein 1 | 1.789027985 |

| | | | |
|---|---|---|---|
| Q64339 | Isg15 | Ubiquitin-like protein ISG15 | 7.109336098 |
| Q91V64 | Isoc1 | Isochorismatase domain-containing protein 1 | 0.571561892 |
| P11688 | Itga5 | Integrin alpha-5 | 1.188437117 |
| P05555 | Itgam | Integrin alpha-M | -0.906795777 |
| P09055 | Itgb1 | Integrin beta-1 | 0.432188492 |
| P11835 | Itgb2 | Integrin beta-2 | -0.620064066 |
| Q6WVG3 | Kctd12 | BTB/POZ domain-containing protein KCTD12 | -0.823786399 |
| Q3U0V1 | Khsrp | Far upstream element-binding protein 2 | -0.263098594 |
| O35344 | Kpna3 | Importin subunit alpha-4 | 0.847554076 |
| P70168 | Kpnb1 | Importin subunit beta-1 | -0.286508168 |
| Q61595 | Ktn1 | Kinectin | 0.795380962 |
| Q9EP89 | Lactb | Serine beta-lactamase-like protein LACTB, mitochondrial | -0.742692529 |
| Q9CQ22 | Lamtor1 | ragulator complex protein LAMTOR1 | -0.292286677 |
| Q9CPY7 | Lap3 | cytosol aminopeptidase | 0.71902176 |
| Q6ZQ58 | Larp1 | La-related protein 1 | 0.810336397 |
| Q6A0A2 | Larp4b | la-related protein 4B | 0.621782125 |
| Q60787 | Lcp2 | lymphocyte cytosolic protein 2 | 0.65494043 |
| Q07797 | Lgals3bp | Galectin-3-binding protein | 1.441137322 |
| Q9JL15 | Lgals8 | Galectin-8 | 0.850816516 |
| O08573 | Lgals9 | Galectin-9 | 1.046917677 |
| O89017 | Lgmn | Legumain | -1.097559382 |
| P37913 | Lig1 | DNA ligase 1 | -1.226853401 |
| Q64281 | Lilrb4 | Leukocyte immunoglobulin-like receptor subfamily B member 4 | 0.802328918 |
| Q99JW4 | Lims1 | LIM and senescent cell antigen-like-containing domain protein 1 | -0.319839174 |
| Q9Z0M5 | Lipa | lysosomal acid lipase/cholesteryl ester hydrolase | -0.973756033 |
| Q9DBH5 | Lman2 | Vesicular integral-membrane protein VIP36 | -0.455017633 |
| P48678 | Lmna | Prelamin-A/C | -0.953010356 |
| P14733 | Lmnb1 | Lamin-B1 | -0.408788533 |
| P11152 | Lpl | lipoprotein lipase | -1.590235882 |
| Q8BFW7 | Lpp | Lipoma-preferred partner homolog | 0.44689958 |
| Q91ZX7 | Lrp1 | prolow-density lipoprotein receptor-related protein 1 | -1.97793801 |
| P62311 | Lsm3 | U6 snRNA-associated Sm-like protein LSm3 | -0.728461937 |
| Q8BLN5 | Lss | lanosterol synthase | -0.857971705 |
| P70202 | Lxn | Latexin | 0.590577196 |
| P0CW03 | Ly6c2 | Lymphocyte antigen 6C2 | 3.006474761 |
| Q60767 | Ly75 | Lymphocyte antigen 75 | 1.294786691 |
| P08905 | Lyz2 | lysozyme c-2 | -2.638138645 |
| Q9CQY5 | Magt1 | Magnesium transporter protein 1 | -0.388523683 |
| Q2TBA3 | Malt1 | Mucosa-associated lymphoid tissue lymphoma translocation protein 1 homolog | 1.715709944 |
| O09159 | Man2b1 | Lysosomal alpha-mannosidase | -1.191095622 |
| O54782 | Man2b2 | Epididymis-specific alpha-mannosidase | -0.971050837 |
| Q8K2I4 | Manba | Beta-mannosidase | -0.578685984 |

| | | | |
|---|---|---|---|
| P31938 | Map2k1 | Dual specificity mitogen-activated protein kinase kinase 1 | 1.093455019 |
| P27546 | Map4 | Microtubule-associated protein 4 | 0.574384105 |
| Q61166 | Mapre1 | Microtubule-associated protein RP/EB family member 1 | 0.539093468 |
| P28667 | Marcksl1 | MARCKS-related protein | 2.918105287 |
| Q05512 | Mark2 | Serine/threonine-protein kinase MARK2 | -0.430908927 |
| Q8K310 | Matr3 | Matrin-3 | -0.354245057 |
| P97310 | Mcm2 | DNA replication licensing factor mcm2 | -1.085579861 |
| P25206 | Mcm3 | DNA replication licensing factor mcm3 | -0.644251862 |
| P49717 | Mcm4 | DNA replication licensing factor MCM4 | -0.826062743 |
| P49718 | Mcm5 | DNA replication licensing factor mcm5 | -0.70038889 |
| Q61881 | Mcm7 | DNA replication licensing factor MCM7 | -1.054497669 |
| P14152 | Mdh1 | Malate dehydrogenase, cytoplasmic | -0.120967767 |
| Q91VH6 | Memo1 | Protein MEMO1 | -1.593968205 |
| P21956 | Mfge8 | Lactadherin | -0.834805106 |
| Q9CQ86 | Mien1 | migration and invasion enhancer 1 | 0.529956251 |
| P34960 | Mmp12 | Macrophage metalloelastase | -1.843566363 |
| Q14CH1 | Mocos | Molybdenum cofactor sulfurase | 1.138693593 |
| P23249 | Mov10 | Putative helicase MOV-10 | 2.202884706 |
| A1L314 | Mpeg1 | Macrophage-expressed gene 1 protein | -0.558282667 |
| Q61830 | Mrc1 | macrophage mannose receptor 1 | -1.774528726 |
| Q9CQF0 | Mrpl11 | 39S ribosomal protein L11, mitochondrial | -0.642489939 |
| Q9D1P0 | Mrpl13 | 39S ribosomal protein L13, mitochondrial | -0.918329186 |
| Q9D1I6 | Mrpl14 | 39S ribosomal protein L14, mitochondrial | -1.056707927 |
| Q9CPR5 | Mrpl15 | 39S ribosomal protein L15, mitochondrial | -0.725436747 |
| Q99N93 | Mrpl16 | 39S ribosomal protein L16, mitochondrial | -0.754468022 |
| Q9D8P4 | Mrpl17 | 39S ribosomal protein L17, mitochondrial | -1.103955254 |
| Q9D338 | Mrpl19 | 39S ribosomal protein L19, mitochondrial | -0.933094012 |
| Q9Z2Q5 | Mrpl40 | 39S ribosomal protein L40, mitochondrial | -1.078632505 |
| Q9CQN7 | Mrpl41 | 39S ribosomal protein L41, mitochondrial | -0.437661291 |
| Q9EQI8 | Mrpl46 | 39S ribosomal protein L46, mitochondrial | -0.804085509 |
| Q9CQ40 | Mrpl49 | 39S ribosomal protein L49, mitochondrial | -0.653655663 |
| Q8VDT9 | Mrpl50 | 39S ribosomal protein L50, mitochondrial | -0.909099772 |
| Q9D125 | Mrps25 | 28S ribosomal protein S25, mitochondrial | -1.539771462 |
| Q9D7N3 | Mrps9 | 28S ribosomal protein S9, mitochondrial | -0.745017208 |
| P26041 | Msn | Moesin | 0.446221744 |
| P00405 | Mtco2 | Cytochrome c oxidase subunit 2 | -0.783573615 |
| Q922D8 | Mthfd1 | C-1-tetrahydrofolate synthase, cytoplasmic | -0.423247801 |
| Q3V3R1 | Mthfd1l | Monofunctional C1-tetrahydrofolate synthase, mitochondrial | -0.422027489 |
| Q9CZU3 | Mtrex | Superkiller viralicidic activity 2-like 2 | -0.778629762 |
| O88441 | Mtx2 | Metaxin-2 | -0.412114845 |
| P16332 | Mut | Methylmalonyl-CoA mutase, mitochondrial | -0.95247997 |
| Q99JF5 | Mvd | Diphosphomevalonate decarboxylase | -0.538612812 |
| Q9EQK5 | Mvp | major vault protein | 0.163809782 |
| Q7TPV4 | Mybbp1a | Myb-binding protein 1A | -0.354630701 |

| | | | |
|---|---|---|---|
| Q3THE2 | Myl12b | Myosin regulatory light chain 12B | -0.207878407 |
| P70248 | Myo1f | Unconventional myosin-If | -0.472709619 |
| Q8BWZ3 | Naa25 | N-alpha-acetyltransferase 25, NatB auxiliary subunit | 0.750302293 |
| Q9D7V9 | Naaa | N-acylethanolamine-hydrolyzing acid amidase | -1.531240379 |
| Q9QWR8 | Naga | alpha-N-acetylgalactosaminidase | -0.447308889 |
| Q9QUK4 | Naip2 | Baculoviral IAP repeat-containing protein 1b | 0.607470608 |
| Q99KQ4 | Nampt | nicotinamide phosphoribosyltransferase | 0.956945115 |
| Q9CWZ7 | Napg | Gamma-soluble NSF attachment protein | 0.339404435 |
| O09043 | Napsa | Napsin-A | -0.859075852 |
| Q8K224 | Nat10 | RNA cytidine acetyltransferase | -0.884599372 |
| Q8K4Z3 | Naxe | NAD(P)H-hydrate epimerase | 0.2059918 |
| Q9CQ49 | Ncbp2 | Nuclear cap-binding protein subunit 2 | 1.184285296 |
| Q09014 | Ncf1 | Neutrophil cytosol factor 1 | -0.828153669 |
| O70145 | Ncf2 | Neutrophil cytosol factor 2 | -0.59174004 |
| Q62433 | Ndrg1 | Protein NDRG1 | -0.569802403 |
| Q9ERS2 | Ndufa13 | NADH dehydrogenase [ubiquinone] 1 alpha subcomplex subunit 13 | -1.190124642 |
| Q9CQ75 | Ndufa2 | NADH dehydrogenase [ubiquinone] 1 alpha subcomplex subunit 2 | -1.315115447 |
| Q9CPP6 | Ndufa5 | NADH dehydrogenase [ubiquinone] 1 alpha subcomplex subunit 5 | -0.555766217 |
| Q9DCJ5 | Ndufa8 | NADH dehydrogenase [ubiquinone] 1 alpha subcomplex subunit 8 | -0.399196963 |
| Q9DCS9 | Ndufb10 | NADH dehydrogenase [ubiquinone] 1 beta subcomplex subunit 10 | -0.254052932 |
| Q9CQZ6 | Ndufb3 | NADH dehydrogenase [ubiquinone] 1 beta subcomplex subunit 3 | -0.355128198 |
| Q9CQC7 | Ndufb4 | NADH dehydrogenase [ubiquinone] 1 beta subcomplex subunit 4 | -0.248293491 |
| Q9CQ54 | Ndufc2 | NADH dehydrogenase [ubiquinone] 1 subunit C2 | -1.177036209 |
| Q91VD9 | Ndufs1 | NADH-ubiquinone oxidoreductase 75 kDa subunit, mitochondrial | -1.227065735 |
| Q91WD5 | Ndufs2 | NADH dehydrogenase [ubiquinone] iron-sulfur protein 2, mitochondrial | -1.286677337 |
| Q9DCT2 | Ndufs3 | NADH dehydrogenase [ubiquinone] iron-sulfur protein 3, mitochondrial | -0.912813316 |
| Q9CXZ1 | Ndufs4 | NADH dehydrogenase [ubiquinone] iron-sulfur protein 4, mitochondrial | -1.259561937 |
| Q99LY9 | Ndufs5 | NADH dehydrogenase [ubiquinone] iron-sulfur protein 5 | -1.622985761 |
| Q91YT0 | Ndufv1 | NADH dehydrogenase [ubiquinone] flavoprotein 1, mitochondrial | -0.839986901 |
| Q9D6J6 | Ndufv2 | NADH dehydrogenase [ubiquinone] flavoprotein 2, mitochondrial | -1.036162325 |
| Q9WTK5 | Nfkb2 | Nuclear factor NF-kappa-B p100 subunit | 1.046450348 |
| O54910 | Nfkbie | NF-kappa-B inhibitor epsilon | 0.419820737 |
| Q9QZ23 | Nfu1 | NFU1 iron-sulfur cluster scaffold homolog, mitochondrial | -0.455508598 |
| Q9CRB2 | Nhp2 | H/ACA ribonucleoprotein complex subunit 2 | -0.680019142 |
| O70131 | Ninj1 | ninjurin-1 | 0.955936631 |
| Q810Q5 | Nmes1 | Normal mucosa of esophagus-specific gene 1 protein | 1.08762926 |

| | | | |
|---|---|---|---|
| O35309 | Nmi | N-myc-interactor | 1.797188572 |
| Q9CQS2 | Nop10 | H/ACA ribonucleoprotein complex subunit 3 | -0.522386802 |
| P29477 | Nos2 | Nitric oxide synthase, inducible | 3.991783684 |
| Q9Z0J0 | Npc2 | Epididymal secretory protein E1 | 0.531955686 |
| Q11011 | Npepps | puromycin-sensitive aminopeptidase | 0.209115213 |
| Q9DCJ9 | Npl | N-acetylneuraminate lyase | 0.54855862 |
| P60670 | Nploc4 | Nuclear protein localization protein 4 homolog | 0.405911406 |
| Q99J45 | Nrbp1 | Nuclear receptor-binding protein | 0.46697242 |
| P46460 | Nsf | Vesicle-fusing ATPase | -0.138034333 |
| Q9D020 | Nt5c3a | Cytosolic 5'-nucleotidase 3A | 3.635709524 |
| Q9D020 | Nt5c3a | Isoform 1 of Cytosolic 5'-nucleotidase 3A | 3.635709524 |
| Q02819 | Nucb1 | Nucleobindin-1 | -0.875374354 |
| P81117 | Nucb2 | Nucleobindin-2 | -0.513956247 |
| Q9CQF3 | Nudt21 | Cleavage and polyadenylation specificity factor subunit 5 | -0.219142505 |
| Q9JKX6 | Nudt5 | ADP-sugar pyrophosphatase | 0.350768725 |
| Q8BVU5 | Nudt9 | ADP-ribose pyrophosphatase, mitochondrial | 0.709993595 |
| Q99P88 | Nup155 | nuclear pore complex protein nup155 | -0.890729237 |
| Q9QY81 | Nup210 | Nuclear pore membrane glycoprotein 210 | -0.70516129 |
| Q8BJ71 | Nup93 | Nuclear pore complex protein Nup93 | -0.369269008 |
| Q6PFD9 | Nup98 | nuclear pore complex protein Nup98-Nup96 | -0.496456503 |
| Q99JX7 | Nxf1 | nuclear RNA export factor 1 | -1.030989673 |
| P11928 | Oas1a | 2'-5'-oligoadenylate synthase 1A | 2.729833373 |
| Q8VI93 | Oas3 | 2'-5'-oligoadenylate synthase 3 | 3.914049119 |
| Q8VI94 | Oasl1 | 2'-5'-oligoadenylate synthase-like protein 1 | 5.203355869 |
| Q99PG2 | Ogfr | opioid growth factor receptor | 1.209876943 |
| Q9CZ30 | Ola1 | obg-like ATPase 1 | 0.118117502 |
| Q78XF5 | Ostc | oligosaccharyltransferase complex subunit OSTC | -0.361542955 |
| Q62422 | Ostf1 | osteoclast-stimulating factor 1 | 0.329644676 |
| Q60715 | P4ha1 | prolyl 4-hydroxylase subunit alpha-1 | 1.146079548 |
| P29341 | Pabpc1 | Polyadenylate-binding protein 1 | 0.366011585 |
| Q9ET54 | Palld | palladin | 0.721214603 |
| O88428 | Papss2 | bifunctional 3'-phosphoadenosine 5'-phosphosulfate synthase 2 | 0.829487981 |
| Q8BZ20 | Parp12 | Poly [ADP-ribose] polymerase 12 | 2.017583381 |
| Q2EMV9 | Parp14 | poly [ADP-ribose] polymerase 14 | 2.406707277 |
| Q8CAS9 | Parp9 | Poly [ADP-ribose] polymerase 9 | 2.078949054 |
| Q9D0B6 | Pbdc1 | Protein PBDC1 | 0.874115629 |
| Q99MN9 | Pccb | propionyl-CoA carboxylase beta chain, mitochondrial | -1.395395938 |
| Q8BH04 | Pck2 | Phosphoenolpyruvate carboxykinase [GTP], mitochondrial | -0.480536784 |
| P17918 | Pcna | proliferating cell nuclear antigen | -0.613596134 |
| Q8BFP9 | Pdk1 | [Pyruvate dehydrogenase (Acetyl-transferring)] kinase isozyme 1, mitochondrial | -0.598288481 |
| Q8K183 | Pdxk | Pyridoxal kinase | -0.43173249 |
| P70296 | Pebp1 | phosphatidylethanolamine-binding protein 1 | -0.534530292 |

| | | | |
|---|---|---|---|
| Q5SUR0 | Pfas | Phosphoribosylformylglycinamidine synthase | -0.383399357 |
| Q9DCD0 | Pgd | 6-phosphogluconate dehydrogenase, decarboxylating | -0.129558259 |
| Q9D0F9 | Pgm1 | Phosphoglucomutase-1 | 0.953555427 |
| Q9JJT9 | Phax | Phosphorylated adapter RNA export protein | -0.41593493 |
| P67778 | Phb | Prohibitin | -0.515460994 |
| E9Q3L2 | Pi4ka | phosphatidylinositol 4-kinase alpha | -0.416365975 |
| Q9EQ32 | Pik3ap1 | Phosphoinositide 3-kinase adapter protein 1 | 1.308917761 |
| Q3TWL2 | Pip4p1 | Type 1 phosphatidylinositol 4,5-bisphosphate 4-phosphatase | -0.924368459 |
| Q8VEB4 | Pla2g15 | Group XV phospholipase A2 | -1.104663909 |
| P51432 | Plcb3 | 1-phosphatidylinositol 4,5-bisphosphate phosphodiesterase beta-3 | 0.833334141 |
| O35405 | Pld3 | Phospholipase D3 | -1.636272379 |
| Q8BG07 | Pld4 | Phospholipase D4 | -1.011386044 |
| Q9JHK5 | Plek | pleckstrin | 0.628026014 |
| Q9ERS5 | Plekha2 | Pleckstrin homology domain-containing family A member 2 | 0.552197838 |
| Q91WB4 | Plekhf2 | Pleckstrin homology domain-containing family F member 2 | 0.813172779 |
| Q8K124 | Plekho2 | pleckstrin homology domain-containing family O member 2 | 0.267905169 |
| P43883 | Plin2 | perilipin-2 | 0.827071443 |
| Q9DBG5 | Plin3 | Perilipin-3 | -0.239090391 |
| Q99JY8 | Plpp3 | Phospholipid phosphatase 3 | 3.572814177 |
| Q60953 | Pml | Protein PML | 1.553879791 |
| Q9CXT8 | Pmpcb | mitochondrial-processing peptidase subunit beta | -0.375182761 |
| P23492 | Pnp | purine nucleoside phosphorylase | 1.286511365 |
| Q9JKP7 | Pole3 | DNA polymerase epsilon subunit 3 | -0.593821146 |
| P08775 | Polr2a | DNA-directed RNA polymerase II subunit RPB1 | -0.950397098 |
| Q80UW8 | Polr2e | DNA-directed RNA polymerases I, II, and III subunit RPABC1 | -0.458133931 |
| P60898 | Polr2i | DNA-directed RNA polymerase II subunit RPB9 | -0.77093566 |
| Q9D0W5 | Ppil1 | Peptidyl-prolyl cis-trans isomerase-like 1 | -1.043793283 |
| Q8BQ30 | Ppp1r18 | Phostensin | 0.308480024 |
| Q60996 | Ppp2r5c | Serine/threonine-protein phosphatase 2A 56 kDa regulatory subunit gamma isoform | -0.832878536 |
| P63328 | Ppp3ca | Serine/threonine-protein phosphatase 2B catalytic subunit alpha isoform | -0.313596239 |
| Q7TMR0 | Prcp | lysosomal Pro-X carboxypeptidase | -0.739009816 |
| P35700 | Prdx1 | peroxiredoxin-1 | 1.859504517 |
| P99029 | Prdx5 | Peroxiredoxin-5, mitochondrial | 0.992899597 |
| O08709 | Prdx6 | Peroxiredoxin-6 | 0.9162617 |
| Q69ZK0 | Prex1 | Phosphatidylinositol 3,4,5-trisphosphate-dependent Rac exchanger 1 protein | 0.843574777 |
| Q99PV0 | Prpf8 | Pre-mRNA-processing-splicing factor 8 | -0.291026417 |
| Q61207 | Psap | Prosaposin | -0.723520939 |
| Q9R1P0 | Psma4 | Proteasome subunit alpha type-4 | 0.282898404 |
| Q9Z2U1 | Psma5 | Proteasome subunit alpha type-5 | 0.370509639 |

| | | | |
|---|---|---|---|
| Q9QUM9 | Psma6 | Proteasome subunit alpha type-6 | 0.215863643 |
| O35955 | Psmb10 | Proteasome subunit beta type-10 | 0.1683119 |
| Q60692 | Psmb6 | Proteasome subunit beta type-6 | -0.561747011 |
| Q8VDM4 | Psmd2 | 26S proteasome non-ATPase regulatory subunit 2 | -0.229279115 |
| P97371 | Psme1 | Proteasome activator complex subunit 1 | 0.909906498 |
| P97372 | Psme2 | proteasome activator complex subunit 2 | 0.752167932 |
| Q8R326 | Pspc1 | Paraspeckle component 1 | 0.20413934 |
| P97814 | Pstpip1 | Proline-serine-threonine phosphatase-interacting protein 1 | 0.685529901 |
| P17225 | Ptbp1 | Polypyrimidine tract-binding protein 1 | -0.415367835 |
| Q14C51 | Ptcd3 | pentatricopeptide repeat domain-containing protein 3, mitochondrial | -0.938996839 |
| Q8BWM0 | Ptges2 | Prostaglandin E synthase 2 | -0.745744684 |
| P22437 | Ptgs1 | Prostaglandin G/H synthase 1 | -0.655314716 |
| P35821 | Ptpn1 | Tyrosine-protein phosphatase non-receptor type 1 | 0.439762921 |
| P35831 | Ptpn12 | Tyrosine-protein phosphatase non-receptor type 12 | 0.490966216 |
| Q8BUM3 | Ptpn7 | Tyrosine-protein phosphatase non-receptor type 7 | -0.698194164 |
| P06800 | Ptprc | Receptor-type tyrosine-protein phosphatase C | -0.457818204 |
| Q8R143 | Pttg1ip | pituitary tumor-transforming gene 1 protein-interacting protein | -1.427942734 |
| Q922Q4 | Pycr2 | Pyrroline-5-carboxylate reductase 2 | -0.453830041 |
| Q9DCC4 | Pycr3 | Pyrroline-5-carboxylate reductase 3 | -0.548526159 |
| Q9ET01 | Pygl | Glycogen phosphorylase, liver form | 0.198472958 |
| Q9D1G1 | Rab1b | ras-related protein Rab-1B | -0.401790243 |
| P35282 | Rab21 | Ras-related protein Rab-21 | 0.32622806 |
| Q921E2 | Rab31 | ras-related protein rab-31 | -0.571589307 |
| Q9CZE3 | Rab32 | Ras-related protein Rab-32 | 0.479741191 |
| Q91ZR1 | Rab4b | Ras-related protein Rab-4B | 0.661493146 |
| Q9CQD1 | Rab5a | Ras-related protein Rab-5A | 0.217679417 |
| P51150 | Rab7a | ras-related protein Rab-7a | -0.281904036 |
| Q91X96 | Rabif | Guanine nucleotide exchange factor MSS4 | -0.688235463 |
| P68040 | Rack1 | Receptor of activated protein C kinase 1 | 0.166706927 |
| P34022 | Ranbp1 | Ran-specific GTPase-activating protein | -0.116304663 |
| Q9D0I9 | Rars | arginine--tRNA ligase, cytoplasmic | 0.257335761 |
| Q6PFQ7 | Rasa4 | Ras GTPase-activating protein 4 | 2.104613036 |
| Q8CB96 | Rassf4 | Ras association domain-containing protein 4 | -0.450606118 |
| B2RY56 | Rbm25 | RNA-binding protein 25 | -0.495757938 |
| O89086 | Rbm3 | RNA-binding protein 3 | -0.559657334 |
| Q6A0D4 | Rftn1 | raftlin | 0.685950911 |
| P58801 | Ripk2 | Receptor-interacting serine/threonine-protein kinase 2 | 1.226510739 |
| Q05921 | Rnasel | 2-5A-dependent ribonuclease | 0.949457161 |
| Q9ET26 | Rnf114 | E3 ubiquitin-protein ligase RNF114 | 1.362309649 |
| E9Q555 | Rnf213 | E3 ubiquitin-protein ligase RNF213 | 2.723697392 |
| Q91VI7 | Rnh1 | Ribonuclease inhibitor | -0.367463125 |

| | | | |
|---|---|---|---|
| Q9CQ71 | Rpa3 | Replication protein A 14 kDa subunit | -1.05828108 |
| Q9JJ80 | Rpf2 | Ribosome production factor 2 homolog | -2.155349583 |
| P35979 | Rpl12 | 60S ribosomal protein L12 | -0.335256286 |
| P47963 | Rpl13 | 60S ribosomal protein L13 | -0.521474378 |
| P19253 | Rpl13a | 60S ribosomal protein L13a | -0.263885922 |
| Q9CR57 | Rpl14 | 60S ribosomal protein L14 | -0.303341321 |
| Q9CZM2 | Rpl15 | 60S ribosomal protein L15 | -0.36034034 |
| Q9CPR4 | Rpl17 | 60S ribosomal protein L17 | -0.327224172 |
| P35980 | Rpl18 | 60S ribosomal protein L18 | -0.37952451 |
| P62717 | Rpl18a | 60S ribosomal protein L18a | -0.72452673 |
| P67984 | Rpl22 | 60S ribosomal protein L22 | -0.188980889 |
| P62830 | Rpl23 | 60S ribosomal protein L23 | -0.198933295 |
| Q8BP67 | Rpl24 | 60S ribosomal protein L24 | -0.363512925 |
| P61255 | Rpl26 | 60S ribosomal protein L26 | -0.309497682 |
| P27659 | Rpl3 | 60S ribosomal protein L3 | -0.223498526 |
| P62889 | Rpl30 | 60S ribosomal protein L30 | -0.322462571 |
| P62892 | Rpl39 | 60S ribosomal protein L39 | -0.733765751 |
| P47911 | Rpl6 | 60S ribosomal protein L6 | -0.370995169 |
| P14148 | Rpl7 | 60S ribosomal protein L7 | -0.359669136 |
| P62918 | Rpl8 | 60S ribosomal protein L8 | -0.418825689 |
| P62281 | Rps11 | 40S ribosomal protein S11 | 0.202038807 |
| P62264 | Rps14 | 40S ribosomal protein S14 | -0.402876203 |
| P62270 | Rps18 | 40S ribosomal protein S18 | -0.359774735 |
| Q9CZX8 | Rps19 | 40S ribosomal protein S19 | -0.324687076 |
| Q9CQR2 | Rps21 | 40S ribosomal protein S21 | -0.439885824 |
| P62852 | Rps25 | 40S ribosomal protein S25 | -0.407882381 |
| P62983 | Rps27a | Ubiquitin-40S ribosomal protein S27a | 0.617612765 |
| P62858 | Rps28 | 40S ribosomal protein S28 | -0.708682254 |
| P97351 | Rps3a | 40S ribosomal protein S3a | -0.124311514 |
| P97461 | Rps5 | 40S ribosomal protein S5 | -0.456881718 |
| P62754 | Rps6 | 40S RIBOSOMAL PROTEIN S6 | -0.336827428 |
| P62082 | Rps7 | 40S ribosomal protein S7 | -0.237958914 |
| P62242 | Rps8 | 40S ribosomal protein S8 | -0.464422246 |
| P14206 | Rpsa | 40S ribosomal protein SA | -0.483935275 |
| P07742 | Rrm1 | Ribonucleoside-diphosphate reductase large subunit | -1.192906497 |
| Q8CBB9 | Rsad2 | Radical S-adenosyl methionine domain-containing protein 2 | 5.450440457 |
| Q99LF4 | Rtcb | tRNA-splicing ligase RtcB homolog | -0.211643368 |
| P60122 | Ruvbl1 | RuvB-like 1 | -0.250019995 |
| Q9WTM5 | Ruvbl2 | RuvB-like 2 | -0.275889101 |
| P31725 | S100a9 | Protein S100-A9 | 0.278117878 |
| Q69Z37 | Samd9l | sterile alpha motif domain-containing protein 9-like | 1.498222802 |
| Q60710 | Samhd1 | deoxynucleoside triphosphate triphosphohydrolase SAMHD1 | 0.835906491 |
| Q9JLI8 | Sart3 | Squamous cell carcinoma antigen recognized by T-cells 3 | 0.298723994 |

| | | | |
|---|---|---|---|
| Q8K352 | Sash3 | SAM and SH3 domain-containing protein 3 | -1.314939064 |
| P70122 | Sbds | Ribosome maturation protein SBDS | 1.688494331 |
| Q9ERN0 | Scamp2 | Secretory carrier-associated membrane protein 2 | -0.527799162 |
| O35609 | Scamp3 | Secretory carrier-associated membrane protein 3 | -0.778837421 |
| Q9CQA3 | Sdhb | Succinate dehydrogenase [ubiquinone] iron-sulfur subunit, mitochondrial | -0.742371657 |
| O08547 | Sec22b | Vesicle-trafficking protein SEC22b | 0.51560659 |
| Q9D662 | Sec23b | protein transport protein Sec23B | 0.283028313 |
| Q3U2P1 | Sec24a | Protein transport protein Sec24A | 0.366081858 |
| Q3UPL0 | Sec31a | Protein transport protein Sec31A | 0.130515673 |
| Q64213 | Sf1 | Splicing factor 1 | -0.872662248 |
| Q99NB9 | Sf3b1 | splicing factor 3B subunit 1 | -0.356829923 |
| Q921M3 | Sf3b3 | Splicing factor 3B subunit 3 | -0.365898394 |
| Q923D4 | Sf3b5 | Splicing factor 3B subunit 5 | -0.306812077 |
| Q8VIJ6 | Sfpq | splicing factor, proline- and glutamine-rich | -0.260086479 |
| Q91V61 | Sfxn3 | Sideroflexin-3 | -0.757162014 |
| Q9JI99 | Sgpp1 | Sphingosine-1-phosphate phosphatase 1 | -0.661859228 |
| Q9JJU8 | Sh3bgrl | SH3 domain-binding glutamic acid-rich-like protein | 0.412873801 |
| Q91VW3 | Sh3bgrl3 | SH3 domain-binding glutamic acid-rich-like protein 3 | -0.561782253 |
| Q9JK48 | Sh3glb1 | Isoform 2 of Endophilin-B1 | 0.598222747 |
| Q8K2Q9 | Shtn1 | Shootin-1 | -0.293701202 |
| Q62230 | Siglec1 | sialoadhesin | 2.648933071 |
| P97797 | Sirpa | tyrosine-protein phosphatase non-receptor type substrate 1 | -0.754665806 |
| Q8VDQ8 | Sirt2 | NAD-dependent protein deacetylase sirtuin-2 | 0.235134857 |
| Q8BHK6 | Slamf7 | SLAM family member 7 | 3.202348356 |
| Q8BPX9 | Slc15a3 | Solute carrier family 15 member 3 | 0.929741936 |
| Q9D6M3 | Slc25a22 | Mitochondrial glutamate carrier 1 | 1.176339419 |
| Q8BMD8 | Slc25a24 | Calcium-binding mitochondrial carrier protein SCaMC-1 | -0.497880433 |
| P10852 | Slc3a2 | Isoform 2 of 4F2 cell-surface antigen heavy chain | 0.484303055 |
| P18581 | Slc7a2 | Isoform 2 of Cationic amino acid transporter 2 | 4.438678261 |
| Q9Z127 | Slc7a5 | large neutral amino acids transporter small subunit 1 | 0.564744257 |
| P70441 | Slc9a3r1 | Na(+)/H(+) exchange regulatory cofactor NHE-RF1 | 0.515573763 |
| Q8CBA2 | Slfn5 | schlafen family member 5 | 2.979475787 |
| Q9D8T7 | Slirp | SRA stem-loop-interacting RNA-binding protein, mitochondrial | -0.889452546 |
| Q3TKT4 | Smarca4 | Isoform 2 of Transcription activator BRG1 | -0.274905468 |
| Q9Z0H3 | Smarcb1 | SWI/SNF-related matrix-associated actin-dependent regulator of chromatin subfamily B member 1 | -0.458940976 |
| Q9CU62 | Smc1a | structural maintenance of chromosomes protein 1a | -0.501309131 |
| Q6P5D8 | Smchd1 | structural maintenance of chromosomes flexible hinge domain-containing protein 1 | -0.384164586 |
| Q9DB90 | Smg9 | protein SMG9 | 0.947281702 |

| | | | |
|---|---|---|---|
| P70158 | Smpdl3a | acid sphingomyelinase-like phosphodiesterase 3a | -1.05484874 |
| P58242 | Smpdl3b | acid sphingomyelinase-like phosphodiesterase 3B | 0.502300681 |
| Q6P4T2 | Snrnp200 | U5 small nuclear ribonucleoprotein 200 kDa helicase | -0.406180996 |
| P57784 | Snrpa1 | U2 small nuclear ribonucleoprotein A' | -0.660587663 |
| P62317 | Snrpd2 | Small nuclear ribonucleoprotein Sm D2 | -0.108919444 |
| Q61263 | Soat1 | sterol O-acyltransferase 1 | -1.422266285 |
| P09671 | Sod2 | Superoxide dismutase [Mn], mitochondrial | 0.967495985 |
| Q8C0J6 | Sowahc | ankyrin repeat domain-containing protein SOWAHC | 1.824933623 |
| O35892 | Sp100 | Nuclear autoantigen Sp-100 | 2.097768433 |
| Q8BVK9 | Sp110 | SP110 nuclear body protein | 2.125617037 |
| Q9JIA7 | Sphk2 | Sphingosine kinase 2 | 0.658304498 |
| Q9JJF9 | Sppl2a | Signal peptide peptidase-like 2A | 0.555577283 |
| Q64337 | Sqstm1 | sequestosome-1 | 2.344128357 |
| Q6P069 | Sri | Sorcin | 0.153628448 |
| Q64674 | Srm | spermidine synthase | -0.369342405 |
| Q9DBG7 | Srpra | signal recognition particle receptor subunit alpha | 0.604441731 |
| Q9R0U0 | Srsf10 | Serine/arginine-rich splicing factor 10 | -1.2661242 |
| Q9EPQ7 | Stard5 | stAR-related lipid transfer protein 5 | 0.857339894 |
| P42225 | Stat1 | signal transducer and activator of transcription 1 | 1.855013592 |
| Q9WVL2 | Stat2 | signal transducer and activator of transcription 2 | 2.813846493 |
| P42227 | Stat3 | Signal transducer and activator of transcription 3 | 0.636424906 |
| Q9Z108 | Stau1 | double-stranded RNA-binding protein Staufen homolog 1 | 0.723318011 |
| O55098 | Stk10 | Serine/threonine-protein kinase 10 | -0.283705953 |
| P54227 | Stmn1 | Stathmin | -1.135627069 |
| P54116 | Stom | erythrocyte band 7 integral membrane protein | -1.610609752 |
| Q99JB2 | Stoml2 | Stomatin-like protein 2, mitochondrial | -0.484176075 |
| O55106 | Strn | striatin | -0.120586634 |
| P46978 | Stt3a | Dolichyl-diphosphooligosaccharide--protein glycosyltransferase subunit STT3A | -0.420836097 |
| O70439 | Stx7 | Syntaxin-7 | -0.554211848 |
| Q8R0F3 | Sumf1 | sulfatase-modifying factor 1 | 0.600581802 |
| Q8BJS4 | Sun2 | SUN domain-containing protein 2 | -0.489562999 |
| P48025 | Syk | Tyrosine-protein kinase SYK | 0.301338767 |
| Q7TMK9 | Syncrip | Isoform 2 of Heterogeneous nuclear ribonucleoprotein Q | -0.174161998 |
| Q7TMK9 | Syncrip | Heterogeneous nuclear ribonucleoprotein Q | -0.174161998 |
| Q8BYC6 | Taok3 | Serine/threonine-protein kinase TAO3 | 0.497272388 |
| P21958 | Tap1 | Antigen peptide transporter 1 | 1.131865889 |
| P36371 | Tap2 | antigen peptide transporter 2 | 1.117096172 |
| Q9R233 | Tapbp | Tapasin | 1.222949846 |
| Q8R3D1 | Tbc1d13 | TBC1 domain family member 13 | 0.408549942 |
| P36423 | Tbxas1 | Thromboxane-A synthase | -0.883901789 |

| | | | |
|---|---|---|---|
| Q9CY27 | Tecr | Very-long-chain enoyl-CoA reductase | -0.268992165 |
| Q62351 | Tfrc | Transferrin receptor protein 1 | -1.447645163 |
| P21981 | Tgm2 | Protein-glutamine gamma-glutamyltransferase 2 | 1.112718406 |
| Q7TMY4 | Thoc7 | THO complex subunit 7 homolog | -0.68606788 |
| P62075 | Timm13 | mitochondrial import inner membrane translocase subunit TIM13 | -0.533511538 |
| Q9WVA2 | Timm8a1 | Mitochondrial import inner membrane translocase subunit Tim8 A | -0.881671639 |
| Q8BH58 | Tiprl | TIP41-like protein | 0.394441323 |
| P26039 | Tln1 | Talin-1 | -0.199798428 |
| Q9QUN7 | Tlr2 | toll-like receptor 2 | 1.815407804 |
| P58021 | Tm9sf2 | Transmembrane 9 superfamily member 2 | -2.883528576 |
| Q9CR02 | Tma16 | translation machinery-associated protein 16 | 1.727651143 |
| Q9D1D4 | Tmed10 | Transmembrane emp24 domain-containing protein 10 | -0.443534835 |
| Q8VC04 | Tmem106a | transmembrane protein 106A | 0.686575059 |
| Q61033 | Tmpo | Lamina-associated polypeptide 2, isoforms alpha/zeta | -0.760811354 |
| Q9DCC8 | Tomm20 | Mitochondrial import receptor subunit TOM20 homolog | -0.724432407 |
| Q921T2 | Tor1aip1 | Torsin-1A-interacting protein 1 | 0.946941193 |
| Q8BYU6 | Tor1aip2 | Torsin-1A-interacting protein 2 | 1.821110288 |
| Q9ER38 | Tor3a | Torsin-3A | 1.719329223 |
| O89023 | Tpp1 | Tripeptidyl-peptidase 1 | -0.775816342 |
| Q91XB0 | Trex1 | Three-prime repair exonuclease 1 | 3.032919935 |
| Q61510 | Trim25 | E3 ubiquitin/ISG15 ligase TRIM25 | 0.727834184 |
| Q62318 | Trim28 | Transcription intermediary factor 1-beta | -0.331078373 |
| G5E870 | Trip12 | E3 ubiquitin-protein ligase TRIP12 | 0.451644378 |
| Q9DCG9 | Trmt112 | Multifunctional methyltransferase subunit TRM112-like protein | -0.885033108 |
| Q3UDE2 | Ttll12 | Tubulin--tyrosine ligase-like protein 12 | -0.468575716 |
| P68372 | Tubb4b | Tubulin beta-4B chain | -0.297680544 |
| Q8BFR5 | Tufm | elongation factor Tu, mitochondrial | -0.511711111 |
| Q91W90 | Txndc5 | Thioredoxin domain-containing protein 5 | -0.509766219 |
| Q9CQ79 | Txndc9 | Thioredoxin domain-containing protein 9 | 0.848373141 |
| Q9JMH6 | Txnrd1 | Thioredoxin reductase 1, cytoplasmic | 0.326918279 |
| Q3TW96 | Uap1l1 | UDP-N-acetylhexosamine pyrophosphorylase-like protein 1 | -0.403379837 |
| Q02053 | Uba1 | Ubiquitin-like modifier-activating enzyme 1 | 0.319179202 |
| P63280 | Ube2i | SUMO-conjugating enzyme ubc9 | -0.18276316 |
| Q9JJZ4 | Ube2j1 | ubiquitin-conjugating enzyme e2 j1 | 0.863665146 |
| Q9QZU9 | Ube2l6 | Ubiquitin/ISG15-conjugating enzyme E2 L6 | 2.684150355 |
| Q8R317 | Ubqln1 | Ubiquilin-1 | -1.206812088 |
| A2AN08 | Ubr4 | Isoform 5 of E3 ubiquitin-protein ligase UBR4 | 0.415901148 |
| P70362 | Ufd1 | Ubiquitin fusion degradation protein 1 homolog | -0.600781261 |
| Q91ZJ5 | Ugp2 | UTP--glucose-1-phosphate uridylyltransferase | 0.258688864 |
| P13439 | Umps | uridine 5'-monophosphate synthase | -0.826283778 |

| | | | |
|---|---|---|---|
| Q9CZ13 | Uqcrc1 | Cytochrome b-c1 complex subunit 1, mitochondrial | -0.575940152 |
| Q9CR68 | Uqcrfs1 | cytochrome b-c1 complex subunit Rieske, mitochondrial | -0.397335526 |
| Q8R5H1 | Usp15 | ubiquitin carboxyl-terminal hydrolase 15 | 0.850103319 |
| P57080 | Usp25 | ubiquitin carboxyl-terminal hydrolase 25 | 1.936384406 |
| Q80U87 | Usp8 | ubiquitin carboxyl-terminal hydrolase 8 | -0.504670133 |
| O70404 | Vamp8 | vesicle-associated membrane protein 8 | -0.262994041 |
| Q01853 | Vcp | Transitional endoplasmic reticulum ATPase | -0.228404693 |
| Q8CDG3 | Vcpip1 | Deubiquitinating protein VCIP135 | 0.917962992 |
| Q8BX70 | Vps13c | Vacuolar protein sorting-associated protein 13C | -0.777959139 |
| P40336 | Vps26a | Isoform 2 of Vacuolar protein sorting-associated protein 26A | -0.302469629 |
| Q8R5L3 | Vps39 | Vam6/Vps39-like protein | -0.494785869 |
| Q9CR26 | Vta1 | Vacuolar protein sorting-associated protein VTA1 homolog | -0.595344916 |
| O88384 | Vti1b | vesicle transport through interaction with t-SNAREs homolog 1B | -0.482038213 |
| Q99KC8 | Vwa5a | von Willebrand factor A domain-containing protein 5A | 0.483198264 |
| P32921 | Wars | Isoform 2 of Tryptophan--tRNA ligase, cytoplasmic | 0.621842178 |
| Q5ND34 | Wdr81 | WD repeat-containing protein 81 | -0.370489829 |
| Q8K1I7 | Wipf1 | WAS/WASL-interacting protein family member 1 | -0.986464824 |
| Q6P1B1 | Xpnpep1 | xaa-Pro aminopeptidase 1 | 0.324390749 |
| Q91WQ3 | Yars | Tyrosine--tRNA ligase, cytoplasmic | 0.215386254 |
| P62960 | Ybx1 | Nuclease-sensitive element-binding protein 1 | 0.273413852 |
| Q9JKB3 | Ybx3 | Y-box-binding protein 3 | -0.504476182 |
| Q9QY24 | Zbp1 | Z-DNA-binding protein 1 | 2.715859027 |
| Q6NZF1 | Zc3h11a | Zinc finger CCCH domain-containing protein 11A | 0.604891163 |
| Q3UPF5 | Zc3hav1 | zinc finger CCCH-type antiviral protein 1 | 0.689284031 |
| Q9D115 | Znf706 | Zinc finger protein 706 | -1.539865555 |
| Q8R151 | Znfx1 | NFX1-type zinc finger-containing protein 1 | 2.149824089 |
| Q71FD5 | Znrf2 | E3 ubiquitin-protein ligase ZNRF2 | 0.618953394 |
| Q8WUR0 | #N/A | Protein C19orf12 homolog | 2.957610527 |
| P10404 | #N/A | MLV-related proviral Env protein | -1.088220099 |

### Supplemental Table 2. Lists of STING interacting proteins

| Accession | Gene name | Description |
|---|---|---|
| P60710 | Actb | Actin, cytoplasmic 1 [OS=Mus musculus] |
| Q922U2 | Krt5 | Keratin, type II cytoskeletal 5 [OS=Mus musculus] |
| P50446 | Krt6a | Keratin, type II cytoskeletal 6A [OS=Mus musculus] |
| Q6IME9 | Krt72 | Keratin, type II cytoskeletal 72 [OS=Mus musculus] |
| Q9Z2K1 | Krt16 | Keratin, type I cytoskeletal 16 [OS=Mus musculus] |
| P05213 | Tuba1b | Tubulin alpha-1B chain [OS=Mus musculus] |
| P21956 | Mfge8 | Lactadherin [OS=Mus musculus] |
| Q8BFR5 | Tufm | Elongation factor Tu, mitochondrial [OS=Mus musculus] |
| Q3UV17 | Krt76 | Keratin, type II cytoskeletal 2 oral [OS=Mus musculus] |
| Q3TBT3 | Tmem173 | Isoform 2 of Stimulator of interferon genes protein [OS=Mus musculus] |
| P20152 | Vim | Vimentin [OS=Mus musculus] |
| P13020 | Gsn | Gelsolin [OS=Mus musculus] |
| Q3TRJ4 | Krt26 | Keratin, type I cytoskeletal 26 [OS=Mus musculus] |
| O35744 | Chil3 | Chitinase-like protein 3 [OS=Mus musculus] |
| Q9QWL7 | Krt17 | Keratin, type I cytoskeletal 17 [OS=Mus musculus] |
| P01027 | C3 | Complement C3 [OS=Mus musculus] |
| P62631 | Eef1a2 | Elongation factor 1-alpha 2 [OS=Mus musculus] |
| E9Q557 | Dsp | Desmoplakin [OS=Mus musculus] |
| P62737 | Acta2 | Actin, aortic smooth muscle [OS=Mus musculus] |
| Q8C669 | Peli1 | E3 ubiquitin-protein ligase pellino homolog 1 [OS=Mus musculus] |
| P01029 | C4b | Complement C4-B [OS=Mus musculus] |
| Q3UHH1 | Zswim8 | Zinc finger SWIM domain-containing protein 8 [OS=Mus musculus] |
| P47856 | Gfpt1 | Glutamine--fructose-6-phosphate aminotransferase [isomerizing] 1 [OS=Mus musculus] |
| P70248 | Myo1f | Unconventional myosin-If [OS=Mus musculus] |
| Q5SUA5 | Myo1g | Unconventional myosin-Ig [OS=Mus musculus] |

| | | |
|---|---|---|
| Q9WTI7 | Myo1c | Unconventional myosin-Ic [OS=Mus musculus] |
| P57780 | Actn4 | Alpha-actinin-4 [OS=Mus musculus] |
| Q9CZU3 | Mtrex | Superkiller viralicidic activity 2-like 2 [OS=Mus musculus] |
| Q9CYA6 | Zcchc8 | Zinc finger CCHC domain-containing protein 8 [OS=Mus musculus] |
| Q9QXS1 | Plec | Plectin [OS=Mus musculus] |
| Q8VDD5 | Myh9 | Myosin-9 [OS=Mus musculus] |
| P62830 | Rpl23 | 60S ribosomal protein L23 [OS=Mus musculus] |
| P01942 | Hba | Hemoglobin subunit alpha [OS=Mus musculus] |
| Q9WV32 | Arpc1b | Actin-related protein 2/3 complex subunit 1B [OS=Mus musculus] |
| Accession | Gene name | Description |
| P62874 | Gnb1 | Guanine nucleotide-binding protein G(I)/G(S)/G(T) subunit beta-1 [OS=Mus musculus] |
| P62908 | Rps3 | 40S ribosomal protein S3 [OS=Mus musculus] |
| Q9CPR4 | Rpl17 | 60S ribosomal protein L17 [OS=Mus musculus] |
| Q9CYL5 | Glipr2 | Golgi-associated plant pathogenesis-related protein 1 [OS=Mus musculus] |
| P47757 | Capzb | Isoform 3 of F-actin-capping protein subunit beta [OS=Mus musculus] |
| P25444 | Rps2 | 40S ribosomal protein S2 [OS=Mus musculus] |
| P60867 | Rps20 | 40S ribosomal protein S20 [OS=Mus musculus] |
| P07356 | Anxa2 | Annexin A2 [OS=Mus musculus] |
| P35980 | Rpl18 | 60S ribosomal protein L18 [OS=Mus musculus] |
| Q6ZWN5 | Rps9 | 40S ribosomal protein S9 [OS=Mus musculus] |
| P35700 | Prdx1 | Peroxiredoxin-1 [OS=Mus musculus] |
| P15331 | Prph | Isoform 3u of Peripherin [OS=Mus musculus] |
| P62702 | Rps4x | 40S ribosomal protein S4, X isoform [OS=Mus musculus] |
| Q8CGP2 | Hist1h2bp | Isoform 2 of Histone H2B type 1-P [OS=Mus musculus] |
| Q60765 | Atf3 | Cyclic AMP-dependent transcription factor ATF-3 [OS=Mus musculus] |
| Q3V132 | Slc25a31 | ADP/ATP translocase 4 [OS=Mus musculus] |
| P58137 | Acot8 | Acyl-coenzyme A thioesterase 8 [OS=Mus musculus] |
| P62267 | Rps23 | 40S ribosomal protein S23 [OS=Mus musculus] |
| P59999 | Arpc4 | Actin-related protein 2/3 complex subunit 4 [OS=Mus musculus] |

| | | |
|---|---|---|
| P14131 | Rps16 | 40S ribosomal protein S16 [OS=Mus musculus] |
| P08752 | Gnai2 | Guanine nucleotide-binding protein G(i) subunit alpha-2 [OS=Mus musculus] |
| Q64444 | Ca4 | Carbonic anhydrase 4 [OS=Mus musculus] |
| Q9DC51 | Gnai3 | Guanine nucleotide-binding protein G(k) subunit alpha [OS=Mus musculus] |
| Q9D8B3 | Chmp4b | Charged multivesicular body protein 4b [OS=Mus musculus] |
| Q8VED5 | Krt79 | Keratin, type II cytoskeletal 79 [OS=Mus musculus] |
| P11928 | Oas1a | 2'-5'-oligoadenylate synthase 1A [OS=Mus musculus] |
| Q9R0N7 | Syt7 | Isoform 4 of Synaptotagmin-7 [OS=Mus musculus] |
| P99024 | Tubb5 | Tubulin beta-5 chain [OS=Mus musculus] |
| P68372 | Tubb4b | Tubulin beta-4B chain [OS=Mus musculus] |
| P68369 | Tuba1a | Tubulin alpha-1A chain [OS=Mus musculus] |
| P68368 | Tuba4a | Tubulin alpha-4A chain [OS=Mus musculus] |
| Q62191 | Trim21 | E3 ubiquitin-protein ligase TRIM21 [OS=Mus musculus] |
| Q9JJZ2 | Tuba8 | Tubulin alpha-8 chain [OS=Mus musculus] |
| Q99JY9 | Actr3 | Actin-related protein 3 [OS=Mus musculus] |
| P54987 | Acod1 | Cis-aconitate decarboxylase [OS=Mus musculus] |
| Accession | Gene name | Description |
| Q8JZX4 | Rbm17 | Splicing factor 45 [OS=Mus musculus] |
| P19973 | Lsp1 | Lymphocyte-specific protein 1 [OS=Mus musculus] |
| P25911 | Lyn | Tyrosine-protein kinase Lyn [OS=Mus musculus] |
| P16951 | Atf2 | Cyclic AMP-dependent transcription factor ATF-2 [OS=Mus musculus] |
| Q9R112 | Sqor | Sulfide:quinone oxidoreductase, mitochondrial [OS=Mus musculus] |
| P60843 | Eif4a1 | Eukaryotic initiation factor 4A-I [OS=Mus musculus] |
| P97793 | Alk | ALK tyrosine kinase receptor [OS=Mus musculus] |
| P61161 | Actr2 | Actin-related protein 2 [OS=Mus musculus] |
| P16460 | Ass1 | Argininosuccinate synthase [OS=Mus musculus] |
| Q925I1 | Atad3 | ATPase family AAA domain-containing protein 3 [OS=Mus musculus] |
| P63017 | Hspa8 | Heat shock cognate 71 kDa protein [OS=Mus musculus] |
| Q64213 | Sf1 | Splicing factor 1 [OS=Mus musculus] |
| Q6NXH9 | Krt73 | Keratin, type II cytoskeletal 73 [OS=Mus musculus] |

| | | |
|---|---|---|
| Q02257 | Jup | Junction plakoglobin [OS=Mus musculus] |
| Q91YQ5 | Rpn1 | Dolichyl-diphosphooligosaccharide--protein glycosyltransferase subunit 1 [OS=Mus musculus] |
| P38647 | Hspa9 | Stress-70 protein, mitochondrial [OS=Mus musculus] |
| Q8BMJ8 | Sp8 | Transcription factor Sp8 [OS=Mus musculus] |
| Q62167 | Ddx3x | ATP-dependent RNA helicase DDX3X [OS=Mus musculus] |
| P14733 | Lmnb1 | Lamin-B1 [OS=Mus musculus] |
| O08638 | Myh11 | Myosin-11 [OS=Mus musculus] |
| Q9WUA3 | Pfkp | ATP-dependent 6-phosphofructokinase, platelet type [OS=Mus musculus] |
| A2BIM8 | Mup18 | major urinary protein 18 [OS=Mus musculus] |
| Q8K1L0 | Creb5 | Cyclic AMP-responsive element-binding protein 5 [OS=Mus musculus] |
| Q9WUM4 | Coro1c | coronin-1C [OS=Mus musculus] |
| Q61233 | Lcp1 | Plastin-2 [OS=Mus musculus] |
| P20029 | Hspa5 | 78 kDa glucose-regulated protein [OS=Mus musculus] |
| Q7TPR4 | Actn1 | Alpha-actinin-1 [OS=Mus musculus] |
| E9Q634 | Myo1e | Unconventional myosin-Ie [OS=Mus musculus] |
| Q9WU78 | Pdcd6ip | Isoform 3 of Programmed cell death 6-interacting protein [OS=Mus musculus] |
| Q6IFX2 | Krt42 | Keratin, type I cytoskeletal 42 [OS=Mus musculus] |
| O55143 | Atp2a2 | Sarcoplasmic/endoplasmic reticulum calcium ATPase 2 [OS=Mus musculus] |
| P11499 | Hsp90ab1 | Heat shock protein HSP 90-beta [OS=Mus musculus] |
| P97449 | Anpep | Aminopeptidase N [OS=Mus musculus] |
| Q9Z331 | Krt6b | Keratin, type II cytoskeletal 6B [OS=Mus musculus] |
| Accession | Gene name | Description |
| O35691 | Pnn | Pinin [OS=Mus musculus] |
| Q9JKF1 | Iqgap1 | Ras GTPase-activating-like protein IQGAP1 [OS=Mus musculus] |
| Q99104 | Myo5a | Unconventional myosin-Va [OS=Mus musculus] |
| Q6R891 | Ppp1r9b | Neurabin-2 [OS=Mus musculus] |
| Q8BTM8 | Flna | Filamin-A [OS=Mus musculus] |
| Q80SU7 | Gvin1 | Interferon-induced very large GTPase 1 [OS=Mus musculus] |
| Q0P678 | Zc3h18 | Zinc finger CCCH domain-containing protein 18 [OS=Mus musculus] |

| | | |
|---|---|---|
| B2RQC6 | Cad | CAD protein [OS=Mus musculus] |
| Q8CH25 | Sltm | SAFB-like transcription modulator [OS=Mus musculus] |
| P62983 | Rps27a | Ubiquitin-40S ribosomal protein S27a [OS=Mus musculus] |
